# The RNA exosome maintains cellular RNA homeostasis by controlling transcript abundance in the brain

**DOI:** 10.1101/2024.10.30.620488

**Authors:** Lauryn A. Higginson, Xingjun Wang, Kevin He, Maggie Torstrick, Minhoo Kim, Bérénice A. Benayoun, Adam MacLean, Guillaume F. Chanfreau, Derrick J. Morton

## Abstract

Intracellular ribonucleases (RNases) are essential in all aspects of RNA metabolism, including maintaining accurate RNA levels. Inherited mutations in genes encoding ubiquitous RNases are associated with human diseases, primarily affecting the nervous system. Recessive mutations in genes encoding an evolutionarily conserved RNase complex, the RNA exosome, lead to syndromic neurodevelopmental disorders characterized by progressive neurodegeneration, such as Pontocerebellar Hypoplasia Type 1b (PCH1b). We establish a CRISPR/Cas9-engineered *Drosophila* model of PCH1b to study cell-type-specific post-transcriptional regulatory functions of the nuclear RNA exosome complex within fly head tissue. Here, we report that pathogenic RNA exosome mutations alter activity of the complex, causing widespread dysregulation of brain-enriched cellular transcriptomes, including rRNA processing defects—resulting in tissue-specific, progressive neurodegenerative effects in flies. These findings provide a comprehensive understanding of RNA exosome function within a developed animal brain and underscore the critical role of post-transcriptional regulatory machinery in maintaining cellular RNA homeostasis within the brain.

## Introduction

Intracellular transcriptome homeostasis relies on the strict regulation of RNA degradation and processing by ribonucleases (RNases) to maintain cell function and integrity (Allmang et al., 1999b; Delan-Forino et al., 2017; Doma and Parker, 2007; Kim et al., 2004; Mitchell et al., 1997). RNases play critical roles in RNA degradation pathways to regulate gene expression through co-/post-transcriptional regulatory mechanisms that finely balance cellular RNA levels (Garneau et al., 2007; Han et al., 2024; Schoenberg and Maquat, 2012; Zinder and Lima, 2017). Thus, RNases ensure the fidelity of RNA production and mediate RNA decay. This precise regulation is especially critical within the nervous system, as evidenced by the expanding list of RNA-mediated neurological disorders caused by defects in essential and ubiquitously expressed RNA decay machinery, which cause pathogenic accumulation of coding and non-coding RNAs (Di Donato et al., 2016; Gerstberger et al., 2014; Linder et al., 2015; Wan et al., 2012). Despite the wide-ranging functions and inherent redundancy of RNases, how disruption of RNase activity leads to neuropathology remains enigmatic.

Causal variants in RNases across the major classes that define their distinctive functions have been implicated in neurological disorders: endonucleases that cut RNA internally are linked to Ataxia with Oculomotor Apraxia Type 2 (AOA2) (Moreira et al., 2004), Aicardi-Goutières Syndrome (AGS) (Cristini et al., 2022; Crow et al., 2006; Rice et al., 2013), and distinct subtypes of Pontocerebellar Hypoplasia (Budde et al., 2008; Namavar et al., 2011; Schaffer et al., 2014); and 3’-5’ RNases, which degrade RNA from the 3’ end are associated with several subtypes of Pontocerebellar Hypoplasia (PCH) (Boczonadi et al., 2014; Burns et al., 2018; Lardelli et al., 2017; Wan et al., 2012), as well as Short stature, Hearing loss and Retinitis Pigmentosa, and distinctive facies (SHRF) syndrome (Di Donato et al., 2016). However, 5’-3’ RNases, which hydrolyze RNA from the 5’ end, have not yet been directly implicated in neurological diseases. Given that defects in endonucleases and 3’-5’ ribonucleases cause neurological disease, it is evident that nervous system development and function are particularly vulnerable to disruptions in RNase activity. The mechanisms underlying this hypersensitivity, which results in neuropathology, remain unknown.

Recessive missense mutations in genes encoding structural subunits of a 3’-5’ RNase complex, the RNA exosome, cause syndromic neurodevelopmental disorders that predominantly manifest as distinct subtypes of Pontocerebellar Hypoplasia (PCH) (Burns et al., 2018; Morton et al., 2018; Rudnik-Schoneborn et al., 2013; Wan et al., 2012). Autosomal recessive PCH is a group of heterogeneous rare neurodevelopmental disorders characterized by progressive neurodegeneration (OMIM #618810). Seventeen different PCH subtypes (PCH1-17) have been reported to date. All subtypes share common brain defects, including cerebellar atrophy and motor neuropathy. RNA exosome-linked PCH accounts for approximately 20% of all known cases of this disorder (van Dijk et al., 2018).

The RNA exosome is a 10-subunit complex composed of 3 Cap structural subunits containing putative S1 and KH RNA binding domains, 6 Core structural subunits containing PH-like domains, and a 3’-5’ catalytically active endo/exoribonuclease subunit, Dis3 (illustrated in **Figure 1A**) (Liu et al., 2006; Mitchell et al., 1997). This structural arrangement creates a central channel that allows RNA substrates to be threaded through the Cap and Core rings for processing and degradation by the basal catalytic subunit, which is primarily Dis3 in the cytoplasm and EXOSC10/Rrp6 in the nucleus (Briggs et al., 1998; Dziembowski et al., 2007; Januszyk and Lima, 2014; Januszyk et al., 2011; Schneider and Tollervey, 2013; Tomecki et al., 2010). The RNA exosome is recruited to specific RNA substrates through distinct nuclear and cytoplasmic adaptor and co-factor proteins (Araki et al., 2001; LaCava et al., 2005; Lubas et al., 2011; Schilders et al., 2005; Schneider and Tollervey, 2013; Vasiljeva and Buratowski, 2006; Zinder and Lima, 2017). These interactions enhance RNA exosome activity, as the complex alone has low intrinsic activity. The RNA exosome is responsible for the turnover of all major classes of RNA (Chekanova et al., 2007; Chen et al., 2001; Flynn et al., 2011; Gudipati et al., 2012; Mitchell et al., 1997; Pefanis et al., 2014; Preker et al., 2008; Schneider et al., 2012). RNA exosome subunits are termed EXOSome Components (EXOSC) in humans and Ribosomal RNA processing proteins (Rrp) in yeast and *Drosophila*. In all organisms evaluated, Cap and Core RNA exosome subunits are essential (Allmang et al., 1999b; Briggs et al., 1998; Hou et al., 2012; Kim et al., 2010; Lim et al., 2013; Mitchell et al., 1997). Studies in budding yeast have assessed the functional consequences of RNA exosome disease-causing amino acid changes in RNA exosome subunits (Fasken et al., 2017; Gillespie et al., 2017); however, these studies provide little insight into how these specific changes could cause tissue-specific consequences. In our previous work, we produced the first multicellular genetic model of PCH1b, a subtype of PCH, in *Drosophila*, revealing an increased requirement for RNA exosome subunit 3 (EXOSC3/Rrp40) in neurons (Morton et al., 2020). In addition, loss-of-function studies in zebrafish indicate that RNA exosome function is crucial for brain development (Wan et al., 2012). The extent to which the functional disruption of the RNA exosome impacts cell-type/tissue-specific gene expression programs and causes pathology within the nervous system requires further investigation.

**Fig. 1:**
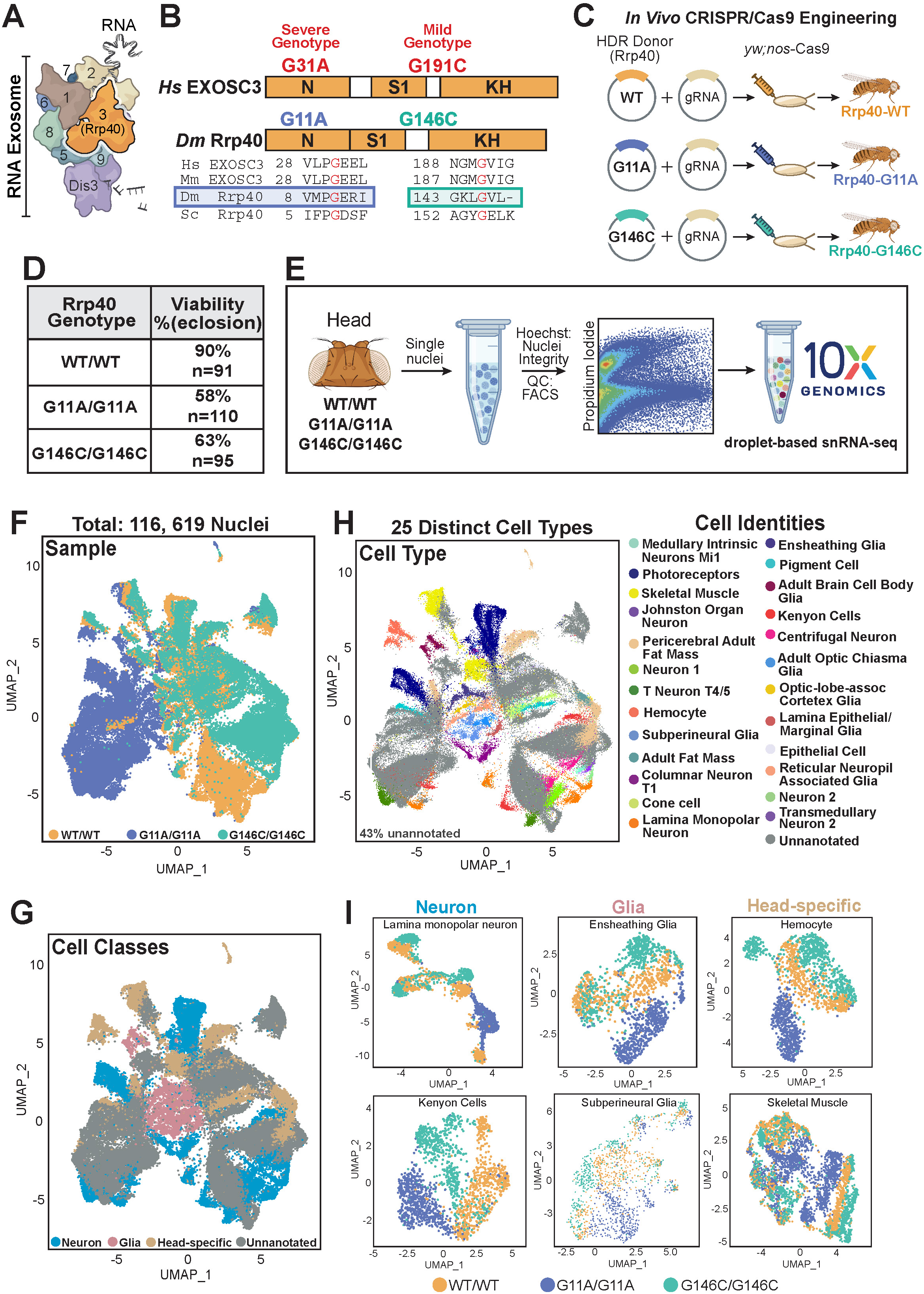
Single-nucleus transcriptome analysis of *Rrp40* mutants reveals distinct cell-type-specific molecular signatures. **(A)** The RNA exosome is an evolutionarily conserved ribonuclease complex composed of structural subunits (EXOSC1-9) (EXOSC4 not visible) and a catalytic subunit (Dis3). **(B)** Pathogenic EXOSC3 (orange, labeled with 3 (termed Rrp40 in *Drosophila*)) variants cause PCH1b. The domain structures of human EXOSC3 and fly Rrp40 proteins highlight the conservation of EXOSC3/Rrp40 throughout evolution and the changes in amino acids in individuals with RNA exosome-linked PCH1b (red). Alignments of the human, mouse, *Drosophila,* and yeast sequences surrounding the amino acids altered in disease are shown. Amino acid changes modeled in this study in RNA exosome cap subunit EXOSC3 are in the N terminal (N) and S1 RNA binding domain. **(C)** Graphical scheme of CRISPR/Cas9 cloning strategy to generate *Rrp40* mutant fly models of EXOSC3-linked PCH1b amino acid substitutions. **(D)** Viability chart of homozygous *Rrp40* wildtype and mutant flies modeling human PCH1b recessive genotypes (WT/WT, G11A/G11A, and G146C/G146C) shown as %eclosion (of expected Mendelian ratio). n= the number of individual flies analyzed. **(E)** Diagram of the snRNA-seq study design. Two pooled biological replicates of 20 newly eclosed (Day 1) adult female flies for each genotype (WT/WT, G11A/G11A, and G146C/G146C). Tissues were dissociated into nuclei suspensions. Flow cytometry was used to determine the quality and quantity of nuclei. Single-nucleus libraries were made using the 10X Genomics Chromium Next GEM Single Cell 3’ kit for approximately 20,000 nuclei per sample. **(F)** 2D Uniform manifold approximation and projection for dimension reduction (UMAP) plots of snRNA-seq data from approximately 116,690 nuclei containing age-matched *Rrp40* mutant (G11A/G11A (blue) or G146C/G146C (teal)) and *Rrp40* wildtype control (WT/WT (yellow)) samples from brain-enriched fly tissue. **(G)** 2D UMAP plots colored by Cell Classes (Neurons (blue), Glia (pink), Head-specific (tan), Unannotated (gray)) integrated across all snRNA-seq datasets and separated by genotype (WT/WT, G11A/G11A, and G146C/G146C) samples. Based on UMAP displayed in 1F. **(H)** 2D UMAP plots colored by 25 Cell Types (specific Cell Identities, right) integrated across all snRNA-seq datasets and separated by genotype (WT/WT, G11A/G11A, and G146C/G146C). Based on UMAP displayed in 1F. **(I)** Broad neural, glial, and head-specific cell classes visualized by 2D UMAP plots on subclustered Cell Classes (Neuron, Glia, and Head-specific) separated and colored by *Rrp40* genotypes (WT/WT (yellow), G11A/G11A (blue), and G146C/G146C (tan).

To investigate the *in vivo*, cell-type-specific functional consequences of neuropathy-causing RNA exosome mutations within the brain, we generated *Drosophila* mutants modeling distinct PCH1b-linked pathogenic variants linked to different severities (severe: G11A/G11A; mild: G146C/G146C) in the fly ortholog of EXOSC3, termed Rrp40 via CRISPR/Cas9-engineering. Our fly model allows for a systematic evaluation of the molecular and organismal consequences of recessive PCH1b genotypes modeled in *Rrp40*. Using high-throughput transcriptomic approaches, confocal brain imaging, and behavioral assays on *Rrp40* mutant flies, we investigate whether cell-type-specific gene expression programs within the brain require unique functions of the RNA exosome complex.

Results from our single-nucleus RNA sequencing (snRNA-seq) and long-read Nanopore-based RNA sequencing datasets revealed that pathogenic *Rrp40* mutations lead to widespread transcriptomic dysregulation, including defective rRNA processing and stabilization of functionally important neuronal transcripts. Intriguingly, snRNA-seq profiling demonstrated that *Rrp40* mutant flies modeling severe (G11A/G11A) or mild (G146C/G146C) PCH1b genotypes exhibited distinct cell-type-specific molecular signatures, indicating that disease-causing mutations modeled in *Rrp40* differentially alter RNA exosome function within the fly brain. We also find that disruption of transcriptome homeostasis in *Rrp40* mutants disproportionately affects specific transcripts, such as *Arc1*. In *Drosophila*, Arc1 is linked to synapse maturation and plasticity (Ashley et al., 2018). We also demonstrate that the RNA exosome tightly regulates *Arc1* mRNA steady-state levels to maintain neuronal homeostasis in a specific tissue within the fly brain, the memory-related mushroom body. Moreover, we report progressive neurodegenerative effects in aged *Rrp40* mutant flies, including decreased lifespan, defects in mushroom body morphology, and short-term memory impairments indicative of central nervous system (CNS) dysfunction that align with *Rrp40* mutant severity. Crucially, we show that pan-neuronal overexpression of *Arc1* in aged fly brains phenocopies mushroom body defects observed in *Rrp40* mutant flies. Together, our results highlight the essential role of the RNA exosome in maintaining neuronal integrity and proper CNS function.

## Results

### Single-nucleus RNA sequencing of brain-enriched tissue from *Rrp40* mutant flies reveals distinct cell-type-specific gene expression signatures

To investigate RNA exosome function within the brain *in vivo*, we generated flies modeling PCH1b disease-causing mutations at the *Rrp40* locus via CRISPR/Cas9 technology. As shown in **Fig. 1A**, the RNA exosome complex is an evolutionarily conserved, multi-subunit complex. The EXOSC3/Rrp40 protein consists of three domains: an N-terminal domain, an S1 putative RNA binding domain, and a K-homology (KH) putative RNA binding domain (**Fig. 1B**). Recessive missense mutations in *EXOSC3* (fly *Rrp40*) encoding single amino acid substitutions cause PCH1b with varying severity (Halevy et al., 2014; Wan et al., 2012; Zanni et al., 2013). The disease-causing amino acid substitutions (severe: G31A (fly G11A) and mild: G191C (fly G146C)) modeled in *Drosophila* in this study are indicated in red above the domain structure and the sequence surrounding these amino acids is shown below. Given that pathogenic mutations in ubiquitously expressed *EXOSC3* cause tissue-specific neurodevelopmental defects that lead to neurodegeneration in humans (Wan et al., 2012), we sought to generate a *Drosophila* model of PCH1b to assess the functional consequences of neuropathy-causing amino acid substitutions. Thus, we engineered flies via CRISPR/Cas9-engineering modeling a wildtype control (Rrp40- WT) and *Rrp40* mutant (Rrp40-G11A or Rrp40-G146C) flies (**Fig. 1C**). *Rrp40* control and mutant flies were engineered as described in Materials and Methods and validated via Sanger sequencing (**S1A Fig**). *Rrp40* mutant flies modeling PCH1b homozygous mutations show reduced viability to adulthood compared to control flies (WT/WT: 91% of expected, n=91, G11A/G11A: 58% of expected, n=110, and G146C/G146C: 63% of expected, n=95) (**Fig. 1D**). Reduced viability to adulthood of severe *Rrp40* mutant (G11A/G11A) compared to control wildtype flies (WT/WT) are in line with our previous study (Morton et al., 2020).

To gain insight into whether cell-type-specific gene expression programs within the brain require distinct functions of the RNA exosome, we profiled nuclei from brain-enriched tissue of newly eclosed (Day 1) engineered wildtype control (WT/WT) and mutant flies (severe: G11A/G11A or mild: G146C/G146C) using droplet-based single-nucleus RNA sequencing (snRNA-seq) (**Fig. 1E**). Notably, transcriptomic analyses employed in this study serves as a read-out for RNA exosome function. Nuclei were profiled from two biological replicates of approximately 20 adult heads for each genotype (WT/WT, G11A/G11A, or G146C/G146C). In total, we successfully sequenced 116,619 high-quality nuclei (**Fig 1E, S1B-C Fig**). A 2D Uniform Manifold Approximation and Projection plot (UMAP) plot analysis of our snRNA-seq results shows G11A/G11A cell clusters occupy distinct regions of the UMAP space with minimal overlap with wildtype control cell clusters (**Fig. 1E**), whereas G146C/G146C cell clusters largely merge, but not completely, with wildtype control cell clusters ( **Fig. 1E**). Moreover, G11A/G11A and G146C/G146C mutant cell clusters occupy distinct regions of the UMAP space with very minimal overlap. These data indicate that distinct PCH1b-linked *Rrp40* alleles (G11A/G11A or G146C/G146C) differentially alter RNA exosome function in post-transcriptional gene regulation, thus resulting in distinct transcriptomic profiles and cellular clustering.

Given the cell-type and tissue-specific pathology caused by RNA exosome defects in the brain, it is critically important to understand the role of the RNA exosome in regulating cellular processes within brain-enriched tissue. Thus, we sought to understand how *Rrp40* mutants affect cell-type-specific transcriptomes within fly head tissue. We first performed dimensionality reduction of all 116, 619 nuclei combined from batch-corrected and integrated *Rrp40* mutants and wildtype control snRNA-seq datasets (**Fig. 1F, S1D Fig**). Next, we subdivided all sequenced nuclei into their respective cell classes (Neuron, Glia, and Head-specific) (**Fig. 1G**) and manually annotated cell types based on previously used marker genes in the fly brain atlas (Croset et al., 2018; Davie et al., 2018; Li, 2021; Li et al., 2022). Accordingly, we annotated 25 known cell types, including eleven neuronal cell types, seven glial subtypes, and seven distinct head-specific cell populations (**Fig. 1H**). In line with published sc/snRNA-seq fly brain datasets, approximately 43% of the cells remained unannotated in control and mutant flies (Croset et al., 2018; Davie et al., 2018; Li et al., 2022). Results from our snRNA-seq dataset allow us to assess how defects within the RNA exosome alter cell-type-specific transcriptomes, as shown in UMAP plot analysis of the 25 annotated cell types within each cell class (Neuron, Glia, Head-specific) from *Rrp40* mutant (G11A/G11A or G146C/G146C) and wildtype control (WT/WT) flies (**Fig. 1I, S2 Fig**). Thus, our high-quality snRNA-seq dataset allows us to assess whether core principles governing nuclear RNA exosome function are general across all cell types or whether cell-type-specific gene expression programs require unique functions of the RNA exosome within fly head tissue.

### Broad transcriptomic dysregulation occurs within brain-enriched tissue in *Rrp40* mutants

We leveraged our snRNA-seq dataset to evaluate how RNA exosome mutations impact cell-type-specific transcriptome expression patterns utilizing brain-enriched tissue from *Rrp40* mutant (severe: G11A/G11A or mild: G146C/G146C) and control (WT/WT) flies. We first analyzed differential expression on neuronal, glial, and head-Higginson *et al*. 8 specific cell populations from *Rrp40* mutants compared to control flies snRNA-seq datasets. Next, we utilized Multidimensional scaling (MDS) to visualize the relationship between *Rrp40* mutants (G11A/G11A or G146C/G146C) and their respective Neuron, Glia, Head-specific, or Unannotated cell populations compared to control *Rrp40* flies (WT/WT) (**Fig 2A**). Each broad cell class (Neuron, Glia, Head-specific, or Unannotated) in *Rrp40* mutant and control flies clustered independently, indicating distinct cell-type-specific molecular signatures between *Rrp40* mutant and control flies. Cell classes within each genotype are clustered together, indicating that differences across samples are global and not specific to one cell type. The MDS plot was generated from two technical replicates representing each genotype (**S1D Fig**). Thus, we can confirm that our data is consistent between both batches for all samples and cell types. Next, to assess how alterations in the RNA exosome directly impact cell-type-specific transcript abundance within the fly head, we performed differential expression analysis on *Rrp40* mutant (G11A/G11A or G146C/G146C) cell classes (Neuron, Glia, Head-specific) compared to control (WT/WT) flies using DESeq2. Interestingly, when comparing both *Rrp40* mutants (G11A/G11A or G146C/G146C) to control flies, each cell class has approximately the same number of differentially expressed transcripts (Neuron: (892), Glia: (789), and Head-specific: (856) in G11A/G11A flies and in G146C/G146C flies (Neuron: (584), Glia: (551), and Head-specific: (608) (**Fig. 2B**). Differential expression analysis of distinct cell subpopulations in each *Rrp40* mutant compared to control flies with at least 100 cells per genotype are represented by volcano plots in **S3-4 Figs**. Subsequently, we assessed differentially expressed transcripts in both *Rrp40* mutant samples compared to controls. A majority of differentially expressed transcripts in flies modeling the severe *Rrp40* allele (G11A/G11A) compared to controls (WT/WT) were increased (59% of total transcripts) in steady-state RNA transcript levels. In flies modeling the mild *Rrp40* allele (G146C/G146C), the number of differentially abundant (increased or decreased) transcripts is approximately the same. Specifically, of the total 2,537 transcripts that change 2-fold or more in the G11A/G11A mutant samples across all three broad cell classes (Neuron, Glia, and Head-specific), 1,502 (59.2%) are increased (FDR<0.05), while 858 are decreased. Whereas, in G146C/G146C mutant samples modeling mild PCH1b disease, 1,743 transcripts change 2-fold or more across all cell classes (Neuron, Glia, and Head-specific), which is significantly less than *Rrp40* severe mutants (G11A/G11A). Specifically, in G146C/G146C mutant samples, 857 (49.2%) transcripts are increased, and 886 (50.8%) transcripts are decreased (**Fig 2B**). Notably, we observe increased transcriptomic dysregulation in G11A/G11A mutants compared to G146C/G146C mutants. These data represent the average of the two pooled biological replicates. Results from these analyses indicate that distinct PCH1b-linked *Rrp40* mutations differentially impact RNA exosome function, which supports observed genotype-phenotype correlations in individuals with PCH1b (Rudnik-Schoneborn et al., 2013).

**Fig. 2:**
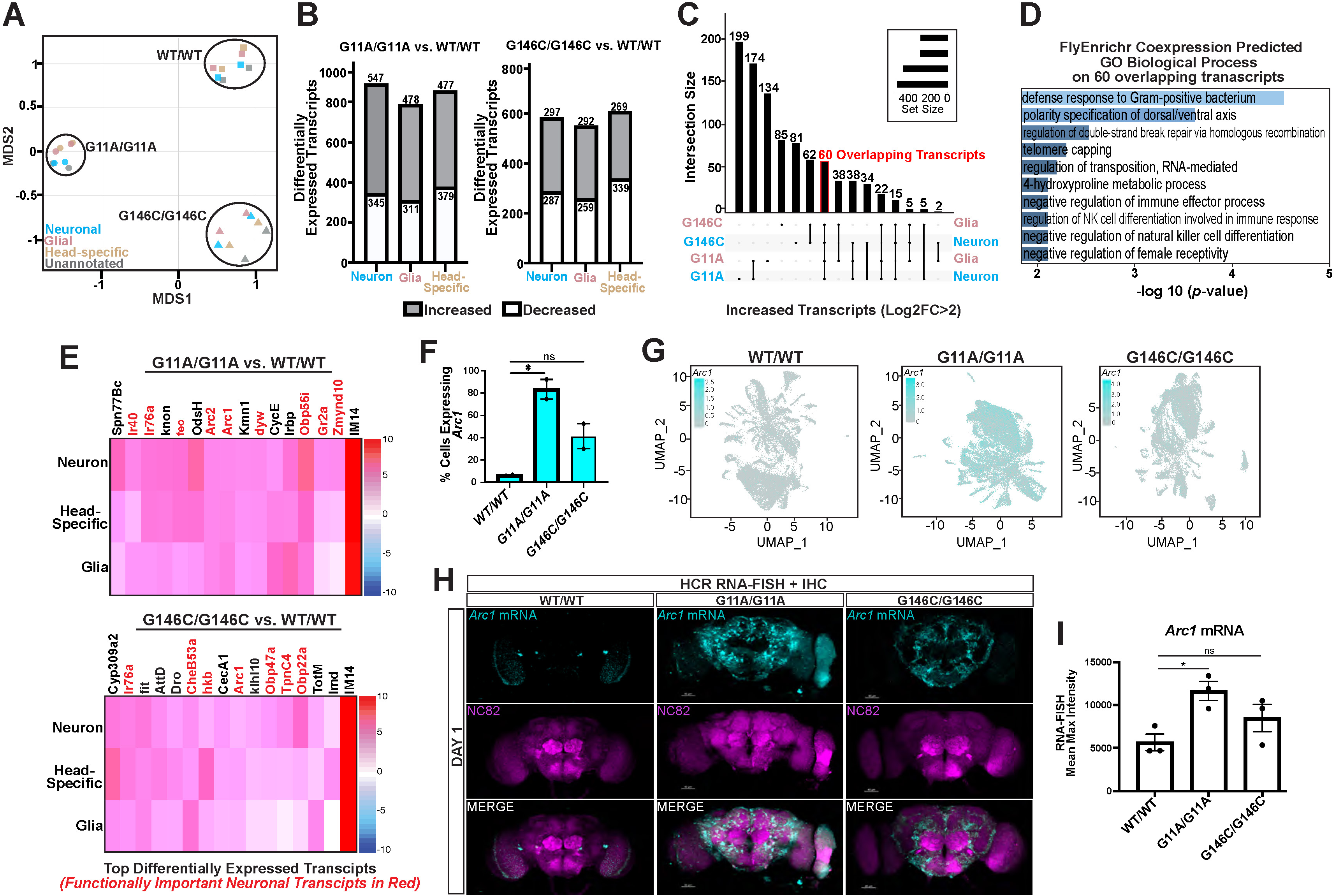
*Rrp40* mutations cause widespread transcriptomic differences across the brain. **(A)** Multidimensional Scaling (MDS) plot. Each point is colored by Cell Class subpopulation (Neuronal (blue), Glial (pink), Head-specific (tan), and Unannotated (gray) and separated by genotype (WT/WT (square), G11A/G11A (circle), and G146C/G146C (triangle). **(B)** Bar graphs representing the number of differentially expressed transcripts increased (gray) (Log2FoldChange>2, FDR<0.05) or decreased (white) (Log2FoldChange<-2, FDR<0.05) separated by genotype compared to control (left, G11A/G11A vs. WT/WT) or (right, G146C/G146C vs. WT/WT)) and colored by Cell Class (Neuron (blue), Glia (pink), and Head-specific (tan)). **(C)** Upset plot of the intersecting differentially expressed transcripts (FDR<0.05) that are increased in Neuron (blue) or Glia (pink) separated by genotype (G11A/G11A or G146/G146C) compared to control (WT/WT). Upset plot highlights 60 overlapping transcripts across each genotype and cell class (neuron or glial) indicated by a red box. **(D)** Bar graph representation of GO Biological Process enriched pathways from FlyEnrichr database (https://maayanlab.cloud/FlyEnrichr/) are shown for the 60 overlapping increased transcripts identified in 2C integrated across snRNA-seq datasets from Neuronal and Glial subpopulations in each *Rrp40* mutant (G11A/G11A or G146C/G146C) as compared to control (WT/WT). The bars shown in blue correspond to significant enrichment (Adj. p-val<0.05). GO terms are ordered from highest p-value significance to lowest. **(E)** Heatmap of the differentially expressed transcripts in (top, G11A/G11A) and (bottom, G146C/G146C) compared to the wildtype control. Functionally important neuronal transcripts are highlighted in red. Normalized Log2FoldChange of differentially expressed transcripts is depicted by a positive 10 (red) gradient to negative - 10 (blue). **(F)** Quantification of the percentage of cells that express *Arc1* in each *Rrp40* mutant (G11A/G11A or G146C/G146C) and wildtype control (WT/WT) from the FeaturePlot displayed in 2G (ns; not statistically significant, *p < 0.05) **(G)** 2D UMAP of FeaturePlot labeling cells that expresses *Arc1* colored in teal in (left, WT/WT) (middle, G11/G11A), and (right, G146C/G146C) cells integrated from 2 pooled biological replicates. The magnitude of *Arc1* expression is depicted by a high-to-low gradient scale (teal to gray) in each sample. **(H)** *Arc1* mRNA quantification (HCR RNA-FISH) coupled with immunohistochemistry (IHC) in whole-mount *Drosophila* brains using confocal microscopy. Simultaneous detection of *Arc1* mRNA (teal) and neuropil marker, nc82/Bruchpilot (Brp), protein (magenta) in Day 1 *Rrp40* fly brains represented as maximum-intensity projections. *Rrp40* flies (Left, WT/WT); (Middle, G11A/G11A); and (Right, G146C/G146C). Top row, *Arc1* mRNA (teal); middle row, NC82 (magenta); and bottom row, merged image (*Arc1* mRNA/NC82) are shown. Scale bars, 50 µm. **(I)** Mean maximum intensity of *Arc1* in *Rrp40* mutants (G11A/G11A or G146C) and *Rrp40* control (WT/WT) from displayed brains in 2H. Asterisks (*) indicate results that are statistically significant at *p-value < 0.05. NS indicates results that show no statistical significance when compared.

Consistent with the primary role of the RNA exosome in RNA decay (Chlebowski et al., 2013; Gudipati et al., 2012), we next focused on transcripts that showed increased expression, as these are likely to be direct targets of the RNA exosome. To determine whether altered RNAs overlap or are distinct between the *Rrp40* mutants (G11A/G11A and G146C/G146C), we generated an UpSet plot to compare brain-specific, differentially expressed transcripts with increased expression across neuronal and glial cell populations (FDR < 0.05, >2-fold change). Interestingly, this analysis revealed substantial overlap in RNAs at the sample level rather than by cell type, suggesting that the RNA exosome targets a common set of transcripts across distinct brain cell types. Among the 1,614 total transcripts that exhibited statistically significant cell-type-specific changes in both *Rrp40* mutants, 38 were altered in glial populations, 34 in neuronal populations, and 60 were altered in both neurons and glia (**Fig. 2C**). Notably, most overlapping transcripts showed a greater magnitude of increase in G11A/G11A compared to G146C/G146C. Additionally, we identified distinct sets of altered RNAs unique to each mutant (G11A/G11A: 507 unique transcripts; G146C/G146C: 228 unique transcripts) (**Fig. 2C**). Together, these data both suggest that RNA exosome activity is more severely compromised in G11A/G11A mutants and that distinct PCH1b variants differentially affect the activity of the complex on specific RNA targets.

Next, we analyzed differentially expressed transcripts from both mutant samples utilizing Gene Ontology (GO) (biological processes) enriched from the 60 overlapping differentially expressed transcripts from each dataset using the FlyEnrichr analysis tool (**Fig 2D**). GO analysis identified 48 statistically significant pathways, including immune response to the bacterium, DNA repair pathways, regulation of transposition, RNA-mediated transposition regulation, and pathways involved in neuronal development. These data suggest that the distinct mutations in RNA exosome genes alter the steady-state levels of a common and distinct set of RNA transcripts critical for cellular homeostasis within the fly brain.

Results presented in **Fig. 2E** show the differentially expressed transcripts in Neuron, Glia, and Head-specific cell clusters from *Rrp40* mutants (G11A/G11A or G146C/G146C) compared to control (WT/WT) and the top 16 differentially expressed transcripts are displayed in the heatmap. Functionally important neuronal transcripts are indicated in red. Of the top 16 differentially abundant transcripts that change 2-fold or more in the G11A/G11A mutant samples, 9 (56%) are functionally important neuronal transcripts. In contrast, 7 of the top 16 transcripts that change 2-fold or more (44%) in the G146C/G146C sample are functionally important neuronal transcripts. The top 16 differentially expressed transcripts in each *Rrp40* mutant compared to control flies are all elevated in each cell class (Neuron, Glia, Head-specific). Of the top differentially abundant transcripts, we identified the neuronal transcript *Arc1*, a key regulator of neuronal function, had the highest percent positive expression across all cell classes (Neuron, Glia, Head-specific) in *Rrp40* mutants. We precisely calculated the percentage of cells expressing *Arc1* in *Rrp40* control and mutant flies: WT/WT: (2,368 of 37,591) 6.3%, G11A/G11A: (34,235 of 41,360) 82.8%, and G146C/G146C (14,839 of 37,568) 39.5% (**Fig. 2F**). These data represent the average of the two pooled biological replicates. Despite increased *Arc1* mRNA levels in mutant flies, Arc1 protein steady-state levels are reduced in *Rrp40* mutants compared to wildtype control flies, as shown by immunoblotting in **S5 Fig**. These data suggest translation efficiency may be compromised in *Rrp40* mutant flies.

To visualize cells expressing *Arc1* mRNA in brain-enriched tissue from *Rrp40* control and mutant flies, we generated a FeaturePlot on a 2D UMAP of all samples independently (**Fig. 2G**). To validate *Arc1* mRNA expression with brain region specificity in *Rrp40* control and mutant flies, we performed Hybridization Chain Reaction (HCR) RNA-FISH coupled with immunohistochemistry (using a presynaptic marker, NC82) on *Rrp40* mutant and wildtype control whole mount fly brains thus allowing visualization of *Arc1* mRNA expression and localization. As shown in **Fig. 2H**, *Arc1* mRNA levels are low and only expressed in the central brain region in wildtype control (WT/WT) brains. In contrast, *Rrp40* mutants (G11A/G11A or G146C/G146C) show both increased *Arc1* mRNA expression and improper distribution as compared to each other and control (WT/WT) flies. Quantification of the mean maximum intensity of these images confirms that *Arc1* mRNA expression in *Rrp40* mutants (G11A/G11A, n=3 or G146C/G146C, n=3) is increased compared to control (WT/WT, n=3) brains (**Fig 2I**). These data demonstrate that PCH1b-linked RNA exosome mutations broadly disrupt transcriptome homeostasis across all cell types within the fly head, leading to the aberrant accumulation and mislocalization of functionally important neuronal transcripts, such as *Arc1*.

### *Rrp40* mutants impair pre-rRNA processing in brain-enriched tissue

The RNA exosome was first identified for its essential role in pre-rRNA processing and is required for *in vivo* 3’ end maturation of 5.8S rRNA (Briggs et al., 1998; Mitchell et al., 1997). To evaluate whether pathogenic RNA exosome mutations alter rRNA biogenesis in *Drosophila* (**Fig. 3A**), as shown in yeast modeling RNA exosome-linked PCH1b mutations (Gillespie et al., 2017), we evaluated rRNA biogenesis in flies modeling PCH1b variants. Thus, we analyzed the effects of G11A/G11A and G146C/G146C mutations on pre-rRNA processing by near-infrared northern blotting (irNorthern) (Miller et al., 2018) on RNA isolated from brain-enriched tissue of newly eclosed (Day 1) flies using a probe hybridizing to 5.8S rRNA. *Rrp40* mutant flies show accumulation of intermediate precursor rRNAs and apparent 3’end extended forms of 5.8S rRNA compared to control flies (**Fig. 3B**). Interestingly, severe *Rrp40* mutants (G11A/G11A) show increased accumulation of precursor and mature 5.8S rRNA compared to mild *Rrp40* mutant (G146C/G146C) analyzed by irNorthern blot (**Fig. 3B**).

**Fig. 3:**
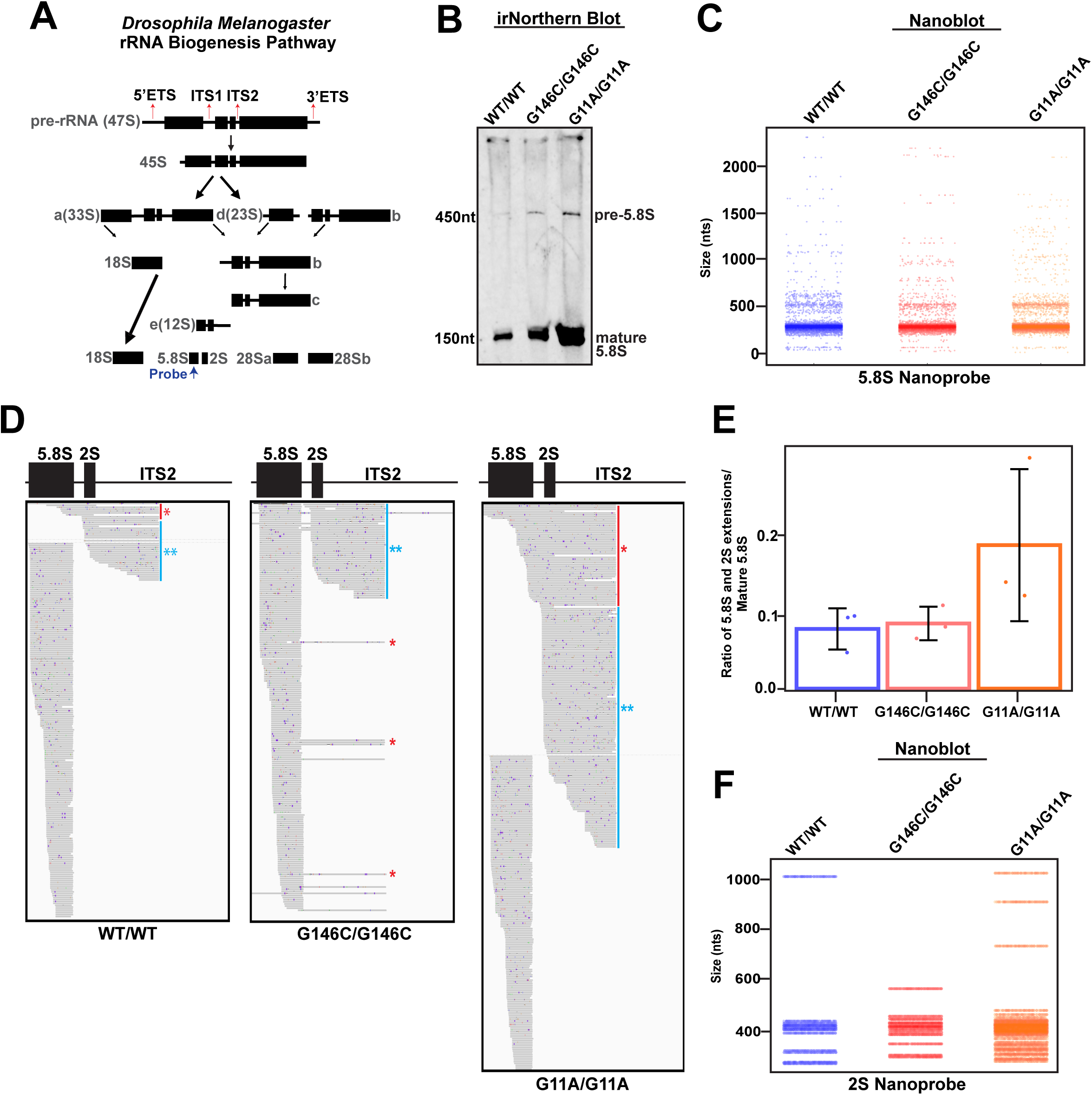
Nanopore long read sequencing in Rrp40 mutant head tissue reveals rRNA biogenesis defects. Overview of *Drosophila melanogaster* ribosomal RNA biogenesis pathway (rRNA) (Gerstberger et al., 2017; Long and Dawid, 1980). The size of rRNA intermediates is indicated in dark gray. Red arrows represent external or internal spacer regions. Interspacer regions are cleaved and excised to produce mature 18S, 5.8S, 2S, 28Sa, and 28Sb rRNA. 5.8S probe (dark blue) was used in near-infrared northern (irNorhern) blots in this study. **(A)** irNorthern blot analysis for 5.8S rRNAs in brain-enriched tissue from *Rrp40* mutants and wildtype control. **(B)** Nanoblot is produced by targeting mature 5.8S rRNA with 5.8S nanoprobe using Nanopore-based RNA-sequencing dataset on *Rrp40* mutants and wildtype controls. Note the similarity of the generated Nanoblot to the irNorthern blot in 3B. **(C)** IGV screenshot of single-track analysis of reads mapping to 5.8S rRNA, 2S rRNA, and ITS2 in *Rrp40* mutants and wildtype control. **(D)** The ratio of 5.8S rRNA and 2S rRNA extensions is quantified in a bar graph for *Rrp40* mutants and wildtype controls. **(E)** Nanoblot is produced by targeting mature 2S rRNA with 2S nanoprobe using the Nanopore-based RNA-Sequencing dataset on *Rrp40* mutants and wildtype controls.

To precisely assess how pathogenic RNA exosome mutations impact rRNA biogenesis in brain-enriched tissue, we applied long-read Nanopore-based RNA-sequencing to *in vitr*o polyadenylated total RNA from 3 independent pooled replicates of newly eclosed (Day 1) *Rrp40* mutant and control fly heads as described in Materials and Methods. From our Nanopore RNA-seq data, we generated northern blot-like images from an R-based package termed Nanoblot, which includes the generation of nanoprobes that can be used to target specific RNAs (DeMario et al., 2023). To recapitulate our irNorthern blot data in 3B, we leveraged our Nanopore RNA-seq dataset to generate a Nanoblot that uses a nanoprobe that targets 5.8S rRNA in *Rrp40* mutant and control fly datasets. In **Fig. 3C**, we show a generated Nanoblot from the Nanopore RNA-seq dataset similar to our irNorthern blot data in 3B. Next, we visualized the rRNA read distribution along the 5.8S, 2S, and internal spacer region 2 (ITS2) rRNA regions by Integrative Genome Viewer (IGV) in *Rrp40* mutant and control flies. Strikingly, as shown in **Fig. 3D**, *Rrp40* mutant flies show a marked increase of 5.8S rRNA species that extended into the ITS2 and defective degradation of the 2S-ITS2 cleavage products compared to *Rrp40* control flies. Intriguingly, severe *Rrp40* mutant (G11A/G11A) had an increased ratio of 5.8S and 2.8S extensions of approximately 0.2 when normalized to mature 5.8S rRNA compared to control flies. In contrast, the ratio of 5.8S-2S extension in the mild *Rrp40* mutant (G146C/G146C) and control flies have similar ratios with a value of approximately 0.1 (**Fig 3E).**

Next, we produced a Nanoblot using nanoprobes targeted to 2S rRNA using our Nanopore RNA-seq dataset. In **Fig. 3F**, our generated Nanoblot shows increased 2S-ITS2 extension of approximately 400nts in *Rrp40* mutants compared to control flies. As control, we utilized our Nanopore RNA-seq dataset to probe 18S rRNA, which does not involve 3’-5’ RNA exosome activity for its maturation. Accordingly, we visualized the rRNA read distribution along the 18S rRNA region in our dataset by IGV and found no 18S rRNA biogenesis defects in *Rrp40* mutant or control flies (**S6 Fig**). We conclude that *Rrp40* mutants modeling PCH1b variants accumulate 3’ extended 5.8S rRNA and show defective degradation of 2S-ITS2 rRNA cleavage product.

### *Rrp40* mutant flies exhibit age-dependent mushroom body γ-lobes defects

To investigate the impact of Rrp40-linked transcriptomic dysregulation (**Fig. 2B**, **Fig. 3A-F, S3-S4 Fig**) on neuronal homeostasis within the fly brain, we assessed the neuroarchitecture of the fly central complex (**Fig. 4A**) and the mushroom body (**Fig. 4B**). The central complex, responsible for higher-order sensory and motor functions (Hulse et al., 2021), consists of the fan-shaped body (FB), protocerebral bridge (PB), ellipsoid body (EB), and antennae lobes (AL) (**Fig. 4A**). The mushroom body, the seat of learning and memory in *Drosophila*, is composed of ∼2,000 intrinsic neurons termed Keyon cells. Kenyon cells (cell bodies) have dendrites that form the mushroom body calyx and axons that extend anteriorly through the peduncle to form the mushroom body lobes (α-, β-, α′-, β′-, and γ-lobes) (de Belle and Heisenberg, 1994; Lin, 2023) (**Fig. 4B**).

**Fig. 4:**
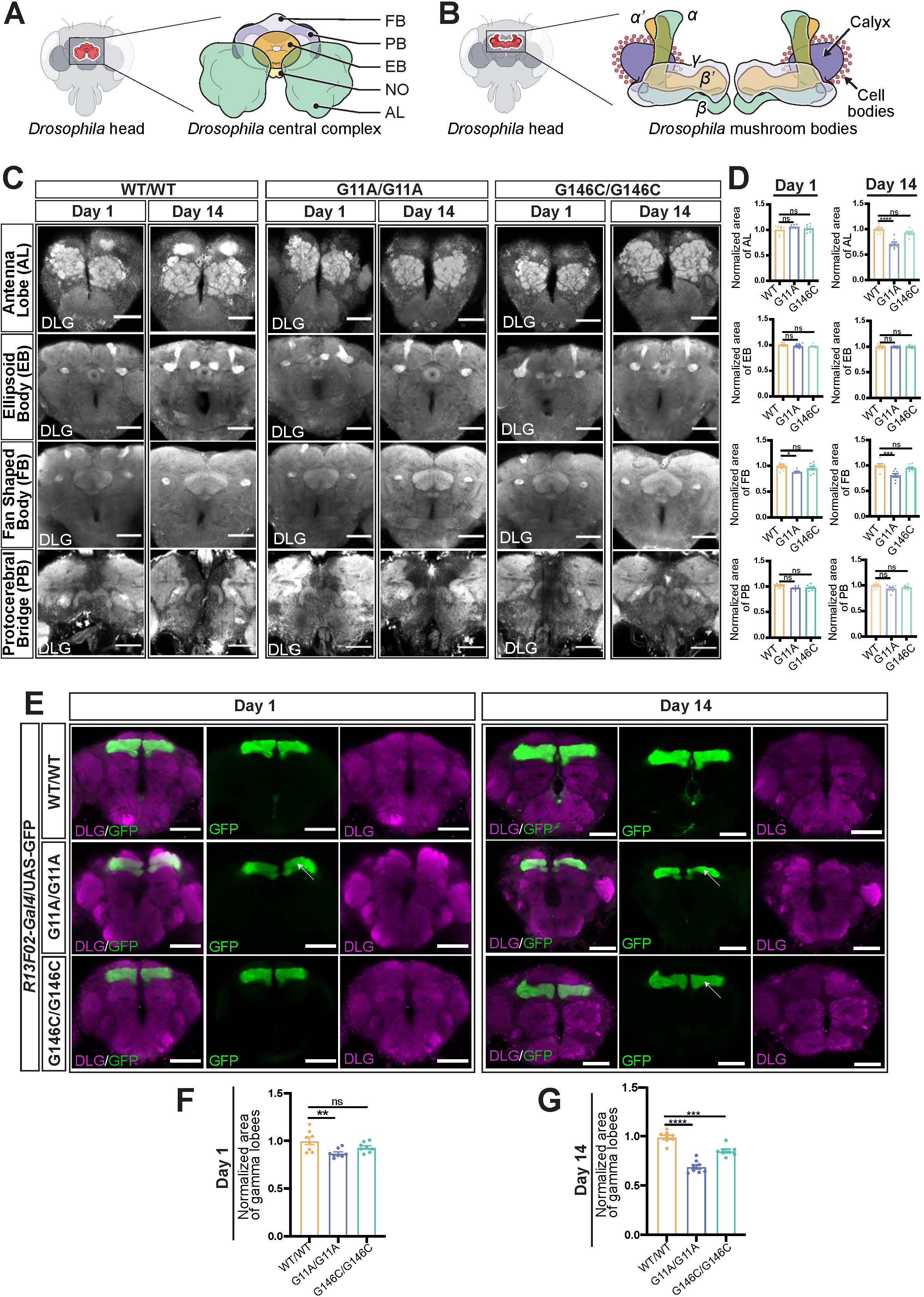
The Rrp40 subunit of the RNA exosome is required for central complex and mushroom body homeostasis within the fly brain. **(A)** Detailed cartoon of the *Drosophila* central complex (left) and the mushroom body (right). The central complex structures highlighted include the fan-shaped body (FB) (light purple), protocerebral bridge (PB) (dark purple, dorsal to FB), ellipsoid body (EB) (orange), and antennae lobes (AL) (green). Depicted mushroom body structure includes vertical alpha (α) (orange) and alpha prime (α′) (light green) neurons, medially projecting beta (β) (orange) and beta prime (β′ (light green), and γ (light purple) neurons and the calyx structure (dark purple), and Kenyon cell bodies (red). **(B)** DLG antibody (anti-DLG) was used to stain *Drosophila* central complex structures on *Rrp40* control (WT/WT) and mutant brains (G11A/G11A or G146C/G146C) on newly eclosed (Day 1) or aged (Day 14) flies using confocal microscopy. Single coronal confocal slices through the center of the brain of central complex structures (top row, (AL); second row (EB); third row, (FB), bottom row, (PB)) from WT/WT (first column), G11A/G11A (middle column), and G146C/G146C (third column) on Day 1 (left) and Day 14 (right) (n=10 per genotype and age) are shown. Scale bar, 100µm. **(C)** Quantification of central complex structures via DLG labeling intensity over time (left column, Day 1 or right column, Day 14) in wildtype controls or *Rrp40* mutant brains. Based on brains displayed in 4B. Asterisks (*) indicate results that are statistically significant at *p-value < 0.05; **p<0.01, ***p<0.001, ****p<0.0001. NS indicates results that show no statistical significance when compared. **(D)** Confocal slices of *Rrp40* mutant and control flies that express GFP under UAS control in combination with mushroom body driver (*R13F02-Gal4) (R13F02-Gal4>UAS-GFP)* from WT/WT (top row), G11A/G11A (middle row), and G146C/G146C (bottom row) flies on Day 1 (left) and Day 14 (right). Brains simultaneously labeled with anti-GFP (mushroom body γ lobe neurons) and anti-DLG (central complex, (magenta) from flies. The first column is a representative merged image (DLG/GFP); the second column is GFP-stained γ lobe neurons (green); and the third column is a DLG-stained central complex (magenta) on newly eclosed (Day 1) and aged (Day 14) flies. Scale bars, 50µm. **(E-F)** Quantification of γ lobes via GFP labeling intensity on Day 1 and Day 14 in wildtype controls or *Rrp40* mutant brains. Based on brains displayed in 4D. Thinned γ -lobes are highlighted by white arrows. Asterisks (*) indicate results that are statistically significant at *p-value < 0.05; **p<0.01, ***p<0.001, ****p<0.0001. NS indicates results that show no statistical significance when compared.

We analyzed the fly central complex in *Rrp40* mutants (G11A/G11A or G146C/G146C) and control flies (WT/WT) by immunostaining for Disc-large (DLG), a post-synaptic neuronal marker, in whole adult fly brains on both Day 1 and Day 14. Confocal microscopy of *Rrp40* control (WT/WT, n=10) and *Rrp40* mutant (G146C/G146C, n=10) reveals no morphological central complex defects on Day 1 (**Fig. 4B**). In contrast, severe *Rrp40* mutant (G11A/G11A, n=10) exhibited a mild reduction in FB size on Day 1 with no apparent gross morphological defects observed in *Rrp40* mutants compared to wildtype controls (**Fig. 4C**). By Day 14, both severe (G11A/G11A, n=10) and mild (G146C/G146C, n=10) *Rrp40* mutants exhibit reduced AL and FB size, with no apparent gross morphological defects observed in mutant flies compared to wildtype controls (**Fig. 4C**). Day 1 and Day 14 results are quantified in **Fig. 4D**. We conclude that *Rrp40* mutants exhibit reductions in AL and FB size but do not show gross morphological central complex abnormalities.

Next, we wanted to assess the role of Rrp40 within specific groups of mushroom body neurons. In our previous work, we reported α- and β-lobe morphogenesis defects in *Rrp40* mutant (G11A/G11A) and *Rrp40^IR^* flies combined with a *Gal4-specific* driver for the mushroom body (*OK107>Gal4*) using the FasII antibody (Morton et al., 2020). Notably, FasII is enriched on α and β axon branches and, to a lesser extent, γ-lobes (Crittenden et al., 1998). Therefore, in our prior study, we could not make confident conclusions about γ-lobe morphogenesis Higginson *et al*. 13 in *Rrp40* mutants. To confirm our prior results and directly assess γ-lobe morphogenesis and neuronal homeostasis in *Rrp40* mutants (G11A/G11A or G146C/G146C), we examined mushroom body structure by genetically combining *Rrp40* control and mutant flies with a UAS/Gal4 system in their respective backgrounds that drives GFP expression in all mushroom body neurons. In addition, we re-examined mushroom body morphology via immunostaining for FasII in *Rrp40* mutant flies.

We first assessed γ-lobe morphogenesis and homeostasis in engineered *Rrp40* control and mutant flies that express GFP from a UAS inducible promoter (*UAS-GFP*) combined with a *Gal4*-specific driver for mushroom body Kenyon cells (*R13F02>Gal4*). *Rrp40* control and mutant animals expressing GFP in mushroom body neurons allow us to directly assess requirements for Rrp40 in distinct groups of mushroom body neurons, including Kenyon cell bodies, dendrites, and axons. Results presented here show confocal sections highlighting mushroom body γ-lobes. First, we analyzed mushroom body γ-lobes via simultaneous staining of GFP and DLG on newly eclosed (Day 1) and aged (Day 14) brains of *Rrp40* mutant and wildtype control flies by confocal microscopy. *Rrp40* mutant (G11A/G11A, n=10) brains exhibit γ-lobe thinning on Day 1 compared to control flies (**Fig. 4E**). Whereas G146C/G146C (n=10) mutants show no γ-lobe defects on Day 1 (**Fig. 4E**). In contrast, both *Rrp40* mutants (G11A/G11A, n=10 or G146C/G146C, n-10) exhibit reduced γ-lobe neuron density by Day 14 compared to control flies as observed by simultaneous staining of GFP and DLG, indicating progressive mushroom body neuronal loss (**Fig. 4E**). Quantification of γ-lobe size in *Rrp40* mutants on Day 1 and Day 14 is shown in **Fig. 4F-G**.

Next, we optically sectioned FasII stained fly brains of *Rrp40* mutant and control flies. Accordingly, we observed robust FasII enrichment on all mushroom body lobes, including FasII-stained γ-lobes albeit reduced in labeling, in newly eclosed and aged control (WT/WT, n=10) and *Rrp40* mutant (G11A/G11A, n=10 or G146C/G146C, n=10) flies (**S7A Fig**). In contrast, FasII staining of aged *Rrp40* mutant flies (G11A/G11A, n=10 or G146C/G146C, n=10) brains on Day 7 and 14 shows reduced labeling of FasII on α and β lobes, revealing α- and β-lobe thinning (**S7A Fig**). In addition, we show no FasII enrichment on γ-lobes in Day 14 severe *Rrp40* mutants (G11A/G11A) compared to control flies, indicating loss of these neurons. Furthermore, we observed reduced FasII enrichment on γ-lobes in aged mild *Rrp40* mutant (G146C/G146C) compared to control flies, suggesting progressive loss of γ-lobe neurons. These data align with our previous report of mushroom body defects in G11A/G11A flies (Morton et al., 2020). Analysis of the mushroom body in G146C/G146C flies modeled Higginson *et al*. 14 in this study serves as the first characterization of this mutant. To further validate the role of Rrp40 in γ-lobe morphogenesis and neuronal homeostasis, we used the *Gal4-*specific mushroom body driver (*R13F02>Gal4*) in combination with RNAi targeting *Rrp40* (BDSC# 62834). Optical sectioning of newly eclosed (Day1) FasII-stained Rrp40-depleted mushroom bodies (*UAS-Rrp40RNAi; R13F02-Gal4*) in flies show no apparent γ-lobe morphogenesis defects compared to controls (*yv;attP40*, BDSC# 36304) (**Fig. S7B**). Whereas, by Day 30, Rrp40-depleted mushroom bodies (*UAS-Rrp40RNAi; R13F02-Gal4*) show reduced γ-lobe density compared to controls (*yv;attP40*, BDSC# 36304) (**S7C Fig**), indicating Rrp40 is critical for mushroom body neuronal homeostasis. From these data, we conclude that Rrp40 is required for proper mushroom body morphogenesis and homeostasis. In sum, *Rrp40* mutant flies exhibit cell-type-specific mushroom body defects but fail to produce gross morphological defects in fly central complex structures.

### *Rrp40* mutant fly brains exhibit decreased neuronal cell populations and mushroom body Kenyon cell death

Given that we observe loss of neuronal homeostasis in *Rrp40* mutant fly mushroom bodies that worsens with age (**Fig. 4D, S7A-C Fig**), we next examined whether we could detect reduced cell proportions in *Rrp40* mutant mushroom body neurons and within head tissue broadly compared to wildtype controls using our snRNA-seq dataset. We used a single-nucleus RNA-seq-specific proportion tool (Miller et al., 2021) to quantify cell type proportions on Day 1 *Rrp40* mutants and wildtype controls. We first analyzed the cell proportions in mushroom body Kenyon cells and broad cell classes (Neuron, Glia, Head-specific) of the fly head in *Rrp40* mutants compared to control flies. We first analyzed cell proportions across cell types within *Rrp40* mutant fly heads compared to wildtype controls. *Rrp40* mutants show significantly decreased neuronal cell populations (>1.25log2FD, FDR<0.05) (**Fig. 5A**). Interestingly, our data do not show significant changes in Head-specific, Glial, and Unannotated cell populations in severe *Rrp40* mutants (G11A/G11A). In contrast, we observe a significant increase in Head-specific cells in mild *Rrp40* mutants (G146C/G146C) without significant changes in Glia or Unannotated cell populations (**Fig. 5A**). These data suggest that RNA exosome function is critical for proper neuronal homeostasis within the fly brain.

**Fig. 5:**
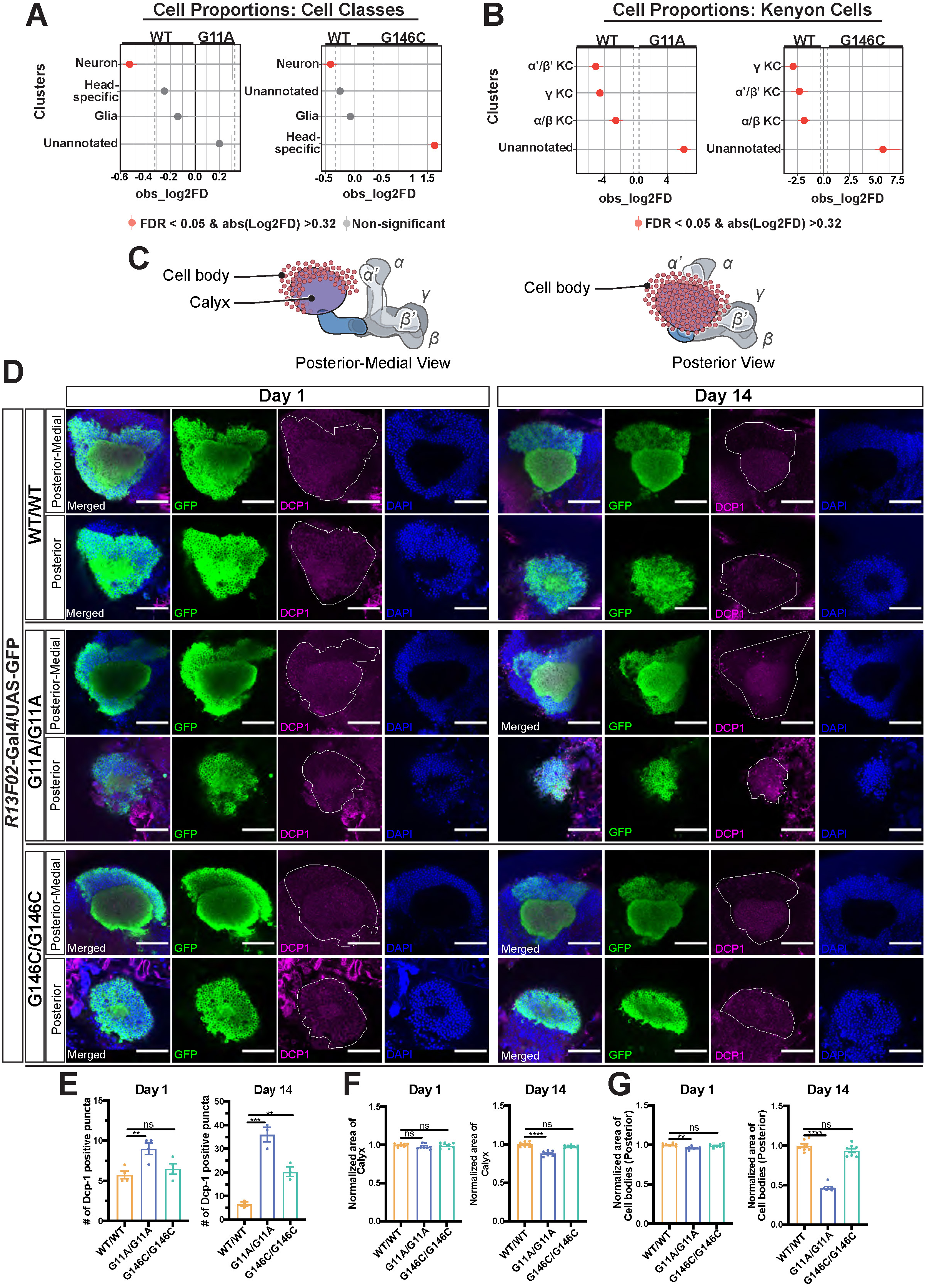
*Rrp40* mutants exhibit neuronal loss within the brain. **(A-B)** Cellular proportion analysis using snRNA-seq dataset on the broad Cell Types (Neuron, Glia, Head-specific, and Unannotated) in *Rrp40* mutants (G11A/G11A or G146C/G146C) compared to control wildtype flies (WT/WT) are shown. Cell proportion analysis of distinct mushroom body Kenyon cell neurons (αβ, α′β′, and γ) in each genotype are shown (B). Cell clusters that are more abundant in the wildtype controls (WT/WT) are represented by dots on the left of the graph, and clusters that are more abundant in the *Rrp40* mutants (G11A/G11A or G146C/G146C) are represented on the right side of the graph. Red dots indicate statistical significance (FDR< 0.05 and absolute (Log2FoldDifference > 0.32), and gray dots indicate non-significant. **(C)** Detailed cartoon of the mushroom body Kenyon cells (red) and calyx (purple) from the posterior-medial and posterior views. **(D)** Representative z-projections in the posterior-medial and posterior view of wildtype controls (WT/WT, top two rows) and *Rrp40* mutants (G11A/G11A, middle two rows or (G146C/G146C, bottom two rows) brains that express GFP under UAS control in combination with mushroom body driver (*R13F02-Gal4) (R13F02-Gal4>UAS-GFP)* labeled with anti-GFP (Cell body and Calyx) (green), anti-DCP-1 (magenta), Cell body and Calyx region of interest outlined in white, and DAPI (blue) on newly enclosed (Day 1, left) and aged (Day 14, right) flies. Scale bar, 100 µm. **(E-G)** Quantification of DCP-1 positive-puncta **(E)**, Calyx size **(F)**, and Cell bodies (posterior view) **(G)** on Day 1 and aged Day 14 *Rrp40* control and mutant flies. Asterisks (*) indicate results that are statistically significant at *p-value < 0.05; **p<0.01, ***p<0.001, ****p<0.0001. NS indicates results that show no statistical significance when compared.

Notably, we observe higher proportions of Unannotated cells in *Rrp40* mutants (G11A/G11A or G146C/G146C) than in control flies (WT/WT) and published sn/scRNA-seq fly head datasets (Davie et al., 2018; Li et al., 2022) in our single-nucleus datasets. These findings suggest that RNA exosome-mediated dysregulation masks the identity of specific cell types. Thus, neuronal loss reported by cell proportion tools in *Rrp40* mutants may be due to the inability to confidently annotate specific neuronal cell types and/or cell death of specific neuronal subtypes, particularly in the severe *Rrp40* mutant (G11A/G11A), which has approximately 1.15 times more unannotated cells (approximately 3,000 cells) than the control. These data indicate that RNA exosome-mediated transcriptomic dysregulation may obscure a higher percentage of cellular identities within the heads of the severe *Rrp40* mutant (G11A/G11A) compared to the mild *Rrp40* mutant (G146C/G146C) and *Rrp40* controls. Next, we compared the cellular proportions of the α, β, α′, β′, and γ Kenyon neuronal cells using a computational cell proportion analysis in R (Miller et al., 2021). In *Rrp40* mutants, our snRNA-seq data show significantly fewer cells for each distinct type of mushroom body neuron (α, β, α′, β′, and γ) compared to control flies (>1.25log2FD, FDR<0.05) (**Fig. 5B**). In addition, in *Rrp40* mutants, our snRNA-seq reveals an increased number of Unannotated Kenyon cells, as observed across cell types within the fly head (**Fig. 5B**), suggesting the RNA exosome-linked RNA dysregulation may also mask the identity of Kenyon cell types. Notably, potential *Rrp40* mutant-mediated masking of cell type identity does not prevent immunohistochemical visualization of the mushroom body neurons in *Rrp40* mutants (**Fig. 4D-G, S7A-B Fig**).

To confirm reduced mushroom body neuron data obtained by cell proportion analyses in **Fig. 5B**, we directly assessed Kenyon cell loss in *Rrp40* mutants compared to control flies via immunohistochemical assays on whole adult fly brains. Thus, we examined cell death in Kenyon cell bodies by immunostaining for the effector caspase Death Caspase 1 (Dcp1) in newly eclosed (Day 1) *Rrp40* control and mutant flies (Day 1) and aged *Rrp40* mutants (Day 14). Kenyon cell bodies are tightly packed around the calyx (Kenyon cell dendrites) (**Fig. 5C**), with each Kenyon cell body dendrite occupying a small proportion of the calyx. Utilizing engineered *Rrp40* control and mutant flies expressing a *Gal4*-specific driver for mushroom bodies combined with a *UAS-GFP* transgene (*Rrp40/Rrp40; R13F02>Gal4/UAS-GFP*), we immunostained whole adult fly brains for Kenyon cell bodies/calyx (Green), Dcp1 (magenta), and DAPI (blue). We imaged from both the posterior and posterior-medial views to gain a comprehensive understanding of Kenyon cell body cell death, given that Kenyon cell bodies wrap around the calyx (**Fig. 5C**). Results demonstrated significant increases in numbers of Dcp1+ Kenyon cell bodies in *Rrp40* mutant brains compared to control flies at Day 1 (**Fig. 5D**), quantified in **Fig. 5E**. Moreover, there was a significant increase in Dcp1+ Kenyon cell bodies in aged severe *Rrp40* mutant (G11A/G11A) compared to mild *Rrp40* mutant (G146C/G146C) and control flies at Day 14 (**Fig. 5D**), quantified in **Fig. 5F**. We conclude that Rrp40 is required for mushroom body morphogenesis and neuronal homeostasis in adult fly brains.

### Pseudo-bulk differential gene expression analysis on subclustered Kenyon cells reveals dysregulation of RNA processing and neuron-specific transcripts in *Rrp40* mutants

Our snRNA-seq analyses and immunohistochemical assays reveal dysregulated transcriptomes and morphogenetic defects in specific mushroom body lobes in *Rrp40* mutants, respectively. Notably, the G11A/G11A mutants exhibit more severe structural abnormalities in mushroom bodies compared to G146C/G146C mutants and control (WT/WT) flies (**Fig. 5A-B**). Therefore, we investigated whether *Rrp40* mutants could be distinguished from each other and control samples based on their mushroom body-specific gene expression profiles.

Therefore, we subclustered Kenyon cells and generated a pseudo-bulk RNA-seq dataset from our snRNA-seq on *Rrp40* mutant (G11A/G11A or G146C/G146C) and control (WT/WT) samples using DESeq2. This analysis enables the examination of cell-type-specific transcriptome changes in neurons of the mushroom body. First, we showed that two technical replicates of *Rrp40* mutant (G11/G11A or G146C/G146C) and control (WT/WT) Kenyon cells clustered separately by genotype using Principal Component Analysis (PCA) ( **Fig. 6A**). These analyses suggest that the pathogenic RNA exosome mutations differentially alter cell-type-specific transcriptomes in the mushroom body. Consistent with results in **Fig. 2B** and **S3-4 Fig** and the role of the RNA exosome in RNA decay (Zinder and Lima, 2017), the majority of the differentially expressed transcripts in *Rrp40* mutants (G11A/G11A or G146C/G146C) were increased by a log2FoldChange of ≥1. Of the 2,554 total transcripts identified, 1455 (57%) in severe *Rrp40* mutant (G11A/G11A) and 1358 (49%) in mild *Rrp40* mutant (G146C/G146C) of a total of 2,357 transcripts were increased by a log2FoldChange >1 compared to control (WT/WT) flies within the mushroom body (**Fig. 6B**).

**Fig. 6:**
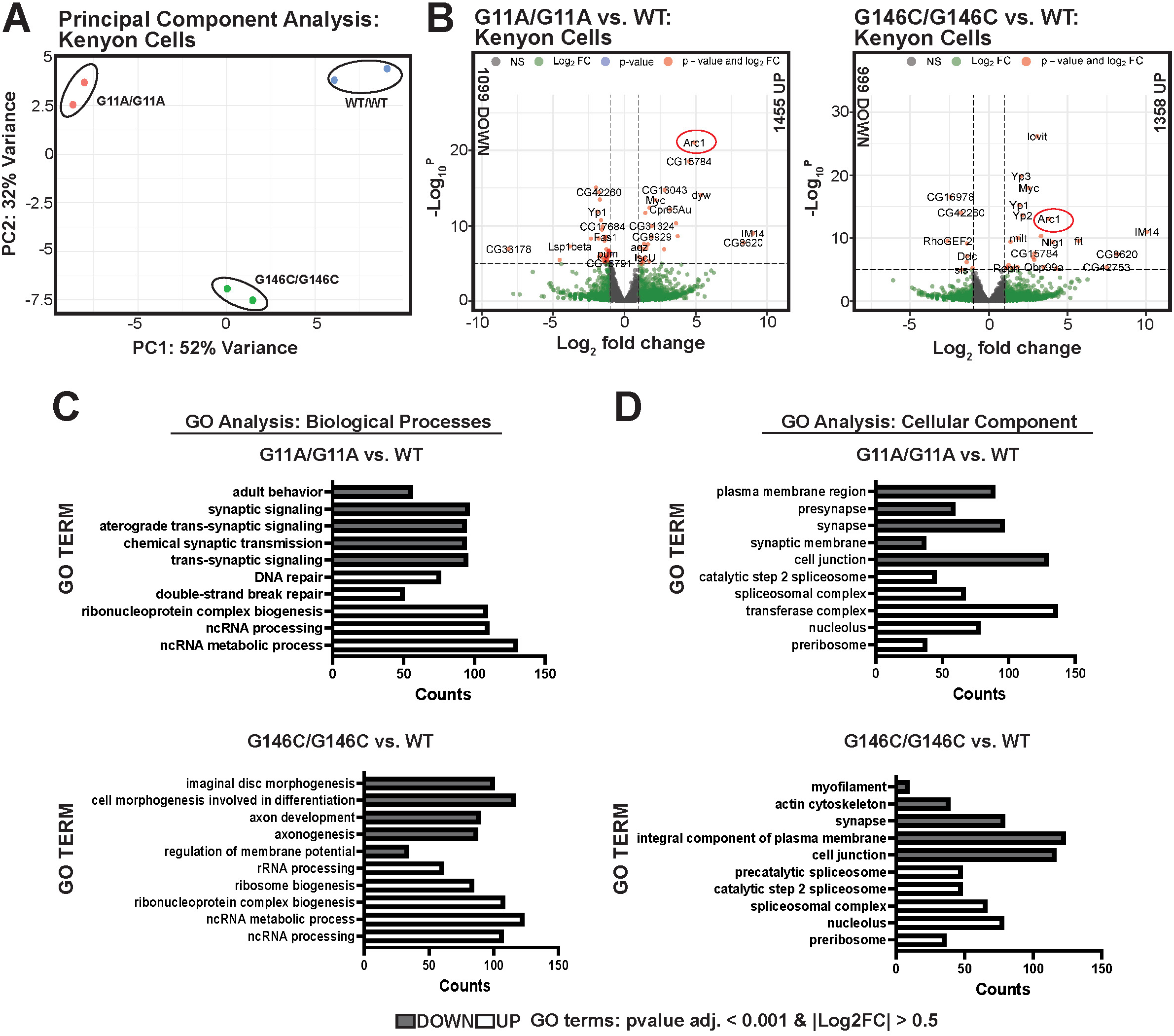
Pseudo-bulked snRNA-seq gene expression analysis in *Rrp40* mutant mushroom body Kenyon cells reveals dysregulated transcripts critical for synaptic transmission and RNA processing. **(A)** Principal component analysis (PCA) plot of two biological replicates of each genotype: WT/WT (blue dots), G11A/G11A (red dots), and G146C/G146C (green dots). PC1 represents 52% of the sample variation, and PC2 represents 32% of the sample variation. **(B)** Volcano plot representing differentially expressed transcripts in G11A/G11A vs. WT/WT (left) and G146C/G146C vs. WT/WT (right). Transcripts (absolute Log2FC > 1) (green), transcripts (p-value < 1e-05) (blue), transcripts (absolute Log2FC > 1 and p-value < 1e-05) (red) , transcripts that meet neither criterion are in gray. **(C-D)** Bar plot of Gene ontology (GO) biological or cellular component **(D)** terms on differentially expressed transcripts (adj. P-value<0.001, absolute Log2FC>0.5) were summarized and selected as described in Materials and Methods. GO analysis processes on genes that are decreased in *Rrp40* mutants vs. control are indicated in gray, and increased transcripts are indicated in white. The top 5 GO terms (increased or decreased differentially expressed transcripts) were plotted for each category.

We next examined the Gene Ontology (GO) enrichment of our samples to identify biological processes (**Fig. 6C**) and cellular components (**Fig. 6D**) that were overrepresented (increased or decreased by a magnitude log2FC of 0.5) in our expression data from *Rrp40* mutants compared to the control. Broadly, biological processes implicated in both *Rrp40* mutants (G11A/G11A or G146C/G146C) for differentially expressed increased transcripts contained RNA processing GO terms, while differentially expressed decreased transcripts contained nervous-system-specific GO terms (**Fig 6C**). Interestingly, from our GO analysis, severe *Rrp40* mutant (G11A/G11A) showed decreased enrichment of synaptic transmission transcripts, with approximately 100 Higginson *et al*. 17 different decreased transcripts implicated in either trans-synaptic signaling or chemical synaptic signaling. In contrast, mild *Rrp40* mutant (G146C/G146C) showed decreased enrichment of axon development, differentiation, and morphogenesis transcripts. Conversely, as expected with disruption of RNA exosome function, we showed that transcripts increased in *Rrp40* mutants (G11A/G11A or G146C/G146C) compared to control are enriched for ncRNA processing, DNA repair, and rRNA processing transcripts (**Fig. 6C**). We also obtained Cellular component analysis (described by their GO term) for *Rrp40* mutants compared to control. Cell component analysis on differentially expressed increased transcripts revealed that both *Rrp40* mutants (G11A/G11A or G146C/G146C) were enriched for spliceosomal complex, nucleolus, and pre-ribosome transcripts (**Fig 6D**). In contrast, differentially expressed decreased transcripts in both *Rrp40* mutants show enrichment for actin cytoskeleton, synapse, and cell junction transcripts (**Fig. 6D**). Thus, pseudo-bulk analysis of subclustered Keyon cells in *Rrp40* mutants reveals altered steady-state transcripts levels important for RNA metabolic processes, axonal neurodevelopment, and synaptic communication.

### *Rrp40* mutants exhibit age-dependent locomotor and taste-learning and memory deficits

The central complex and mushroom body are key regions of the fly brain linked to advanced neuronal functions, including locomotion and learning and memory (Martin et al., 1998; McBride et al., 1999; Strauss, 2002). Defects observed in the central complex and mushroom body in *Rrp40* mutant flies (**Fig. 4A-B, S7A-B Fig**) prompted us to investigate the role of Rrp40 on both the lifespan and functionality of the *Drosophila* central nervous system (CNS). Kaplan-Meier plots of *Rrp40* mutant (G11A/G11A or G146C/G146C) adult survivors revealed significantly reduced lifespans within the 80-day testing period compared to *Rrp40* control flies (WT/WT). Specifically, severe *Rrp40* mutant (G11A/G11A) flies died by Day 22, and mild *Rrp40* mutant (G146C/G146C) flies survived until Day 40 (**Fig 7A**).

**Fig. 7:**
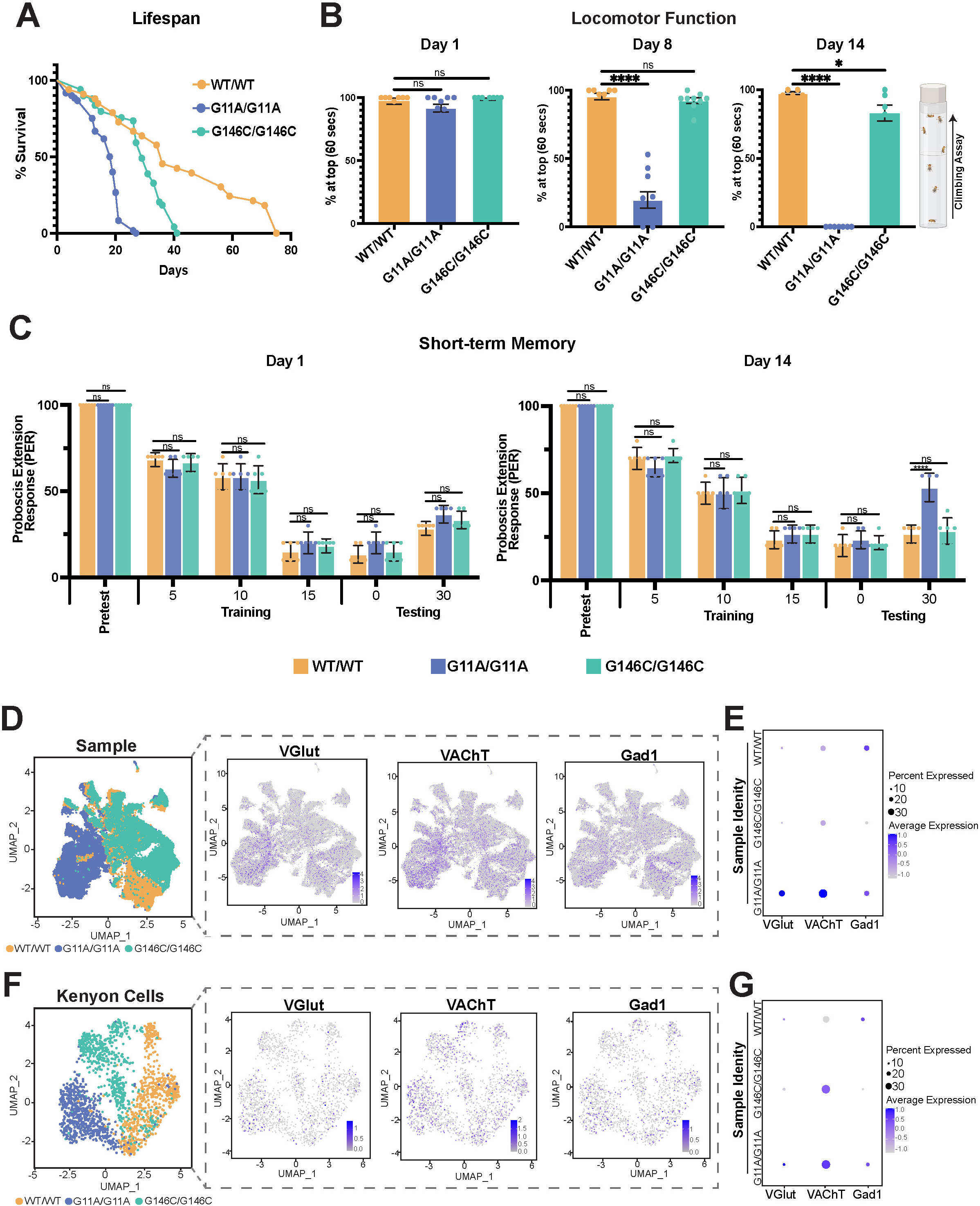
*Rrp40* mutants show a range of organismal and short-term taste memory defects. **(A)** A Kaplan-Meier analysis shows the % survival of *Rrp40* mutant flies with the indicated genotypes: (WT/WT (tan), G11A/G11A (blue), and G146C/G146C (green) (n=100, per genotype). **(B)** Locomotor activity was assessed using a negative geotaxis assay described in Materials and Methods (right, cartoon depiction of climbing assay). Data are presented as the average percentage of flies of the indicated genotypes that reach the top of a cylinder after 60 seconds across all trials. Groups of age-matched (Day 1, Day 8, and Day 14) flies (average n=82, cohorts of 9-12 flies) were tested for at least three independent trials per genotype. Results are represented as mean ± SEM (ns p>0.05,*p<0.05, ****p<0.0001 vs. the WT group in an unpaired two-tailed *t*-test) **(C)** Newly eclosed (left, Day 1) and aged flies (right, Day 14) were subjected to a short-term aversive memory taste assay. Memory assay was performed as described in Materials and Methods. Data represented in bar graphs for *Rrp40* control (WT/WT) and *Rrp40* mutant (G11A/G11 or G146C/G146C) flies. (n=10, per replicate and genotype). All flies were food-deprived for 24 hr before the start of the experimentation. Proboscis-extension rates (PER) are mean ± SEM, 2-way ANOVA test: *p < 0.05, **p < 0.01, ***p < 0.001.

To evaluate the functionality of the *Drosophila* CNS, we conducted negative geotaxis locomotor assays on both *Rrp40* mutant and control flies. *Rrp40* mutants (G11A/G11A or G146C/G146C) demonstrated a progressive decline in locomotor activity with age (**Fig. 7B**). Notably, the more severe G11A/G11A mutants exhibited more significant locomotor impairments compared to the mild G146C/G146C mutants. By Day 14, G11A/G11A mutants could not perform locomotor assays, which aligned with our previous findings (Morton et al., 2020). To investigate whether associative memory formation in the mushroom body is compromised in *Rrp40* mutants, we conducted a short-term memory taste assay as described in the Materials and Methods. Aged Higginson *et al*. 18 severe *Rrp40* mutants (G11A/G11A) displayed taste memory deficits, with 50% of flies responding to the tastant (sucrose) on Day 14. In contrast, mild *Rrp40* mutants (G146C/G146C) and control flies (WT/WT) exhibited similar responses, with 25% responding to sucrose (tastant) **(Fig. 7C)**. These results indicate that mushroom body neuronal defects and loss in G11A/G11A mutants is sufficient to disrupt associative learning and memory functions in flies.

Given the learning and memory deficits observed in *Rrp40* mutant (G11A/G11A) and the GO enrichment analysis revealing significant enrichment of transcripts related to synaptic signaling and transmission in both G11A/G11A and G146C/G146C mutant flies (**Fig. 6C-D)**, we next profiled excitatory and inhibitory neurotransmitters across all cell populations, including mushroom body neurons, using our snRNA-seq dataset. First, we used known markers from publicly available datasets (Davie et al., 2018; Kolodziejczyk et al., 2008) to annotate glutamatergic (VGlut) and cholinergic (VAChT) synaptic transporters and GABAergic (Gad1)+ neurons in *Rrp40* mutants and control flies, analyzing all head cell populations and subclustered mushroom body Kenyon cells. Interestingly, in all cell populations within the fly head, we observed increased expression of VAChT with a normalized average expression of 3.80 in severe *Rrp40* mutant (G11A/G11A), with approximately 33% of head cells expressing VAChT, compared to mild *Rrp40* mutant (G146C/G146C) and wildtype control (WT/WT) flies, where VAChT expression was significantly lower, with an expression of 2.17 and 2.26 in approximately 20% and 21% of head cells, respectively. (**Fig. 7D-E**). Next, we examined VAChT, VGlut, and Gad1 expression in Kenyon cells using our snRNA-seq dataset. We found that VAChT expression was elevated in approximately 40% of Kenyon cells in *Rrp40* mutants, with a normalized average VAChT expression of .59 in G11A/G11A mutants and a normalized VAChT average expression of .55 in G146C/G146C mutants compared to control flies with a normalized VAChT average expression of 0.37 (**Fig. 7F-G**). In contrast, VGlut and Gad1+ cell percentages remained consistent across mutants and controls (**Fig. 7F-G**). *Drosophila* Kenyon cells require accurate VAChT expression for proper function (Barnstedt et al., 2016). Thus, these results are particularly intriguing given that our snRNA-seq dataset shows a reduction in Kenyon cell populations in *Rrp40* mutant heads compared to control flies (**Fig. 4B**). Increased VAChT in *Rrp40* mutants suggests that increased cholinergic signaling may be compensating for Kenyon cell loss. These findings indicate that Rrp40 is essential for maintaining proper cholinergic function in the fly brain.

### Overexpression of *Drosophila Arc1* is sufficient to produce mushroom body defects in aged flies

The integrative survey of cell-type-specific transcriptome changes underpinning each *Rrp40* mutant fly coupled with *in vivo* phenotypes reported here enables us to screen and validate whether specific dysregulated transcripts, such as *Arc1*, contribute to neuronal-specific phenotypes. *Arc1* mRNA levels are disproportionately increased in *Rrp40* mutants (**Figs. 2E-I**). To investigate whether the synaptic regulator *Arc1* influences mushroom body morphogenesis and/or homeostasis *in vivo*, we used the UAS/Gal4 system in flies, employing a commonly used pan-neuronal driver (*elav>Gal4*) (Robinow and White, 1988) to overexpress *Arc1* from a UAS promotor (*UAS-Arc1*). We then examined steady-state Arc1 protein levels in control flies (*w^1118^*, *elav>Gal4*, or *UAS-Arc1*) and in flies expressing *Arc1* (*UAS-Arc1*) driven by pan-neuronal driver (*elav>Gal4*) via immunoblotting. Immunoblots on head tissue from flies expressing *Arc1* with the pan-neuronal driver (*elav>Gal4; UAS-Arc1*) using validated Arc1 antibody (Ashley et al., 2018) show increased steady-state protein levels of both multimeric and monomeric Arc1 species (**Fig. 8A**). Arc monomers form retrovirus-like capsids that encapsulate *Arc* mRNA, allowing this RNA is transferred across synapses via extracellular vesicles (Ashley et al., 2018; Pastuzyn et al., 2018). Studies in *Drosophila* suggest that synaptic plasticity depends on the trans-synaptic transfer of *Arc1* (Ashley et al., 2018).

**Fig. 8:**
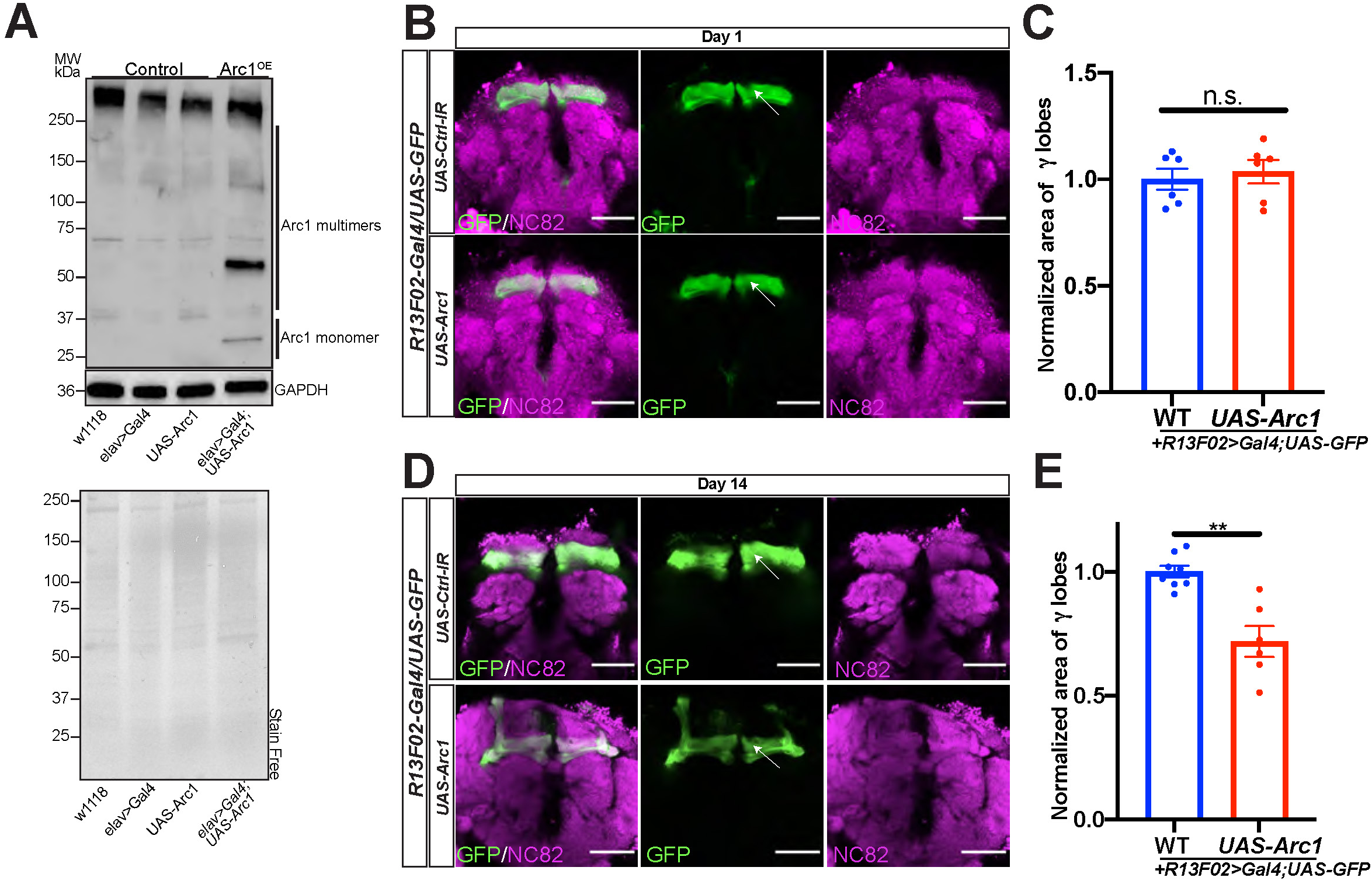
Overexpression of *Arc1* in aged flies exhibit mushroom body defects. **(A)** Western blot on lysates prepared from heads on control flies (w^1118^, *elav>Gal4*, UAS-*Arc1*^OE^) and *elav*-Gal4>UAS>*Arc1*^OE^ were resolved by SDS-PAGE and analyzed by immunoblotting with antibodies to detect Arc1 monomeric and multimeric species. Both GAPDH and Stain Free, as a measure of total protein, serve as loading controls. **(B)** Confocal slices of *Arc1^OE^* (UAS-Arc1, BSDC #37532) and control flies (R13F02>Gal4/UAS/GFP; Control-IR) (Control IR, BDSC #36303) that express GFP under UAS control in combination with mushroom body driver (*R13F02-Gal4) (R13F02-Gal4>UAS-GFP)* on Day 1. Brains were simultaneously labeled with anti-GFP (mushroom body γ-lobe neurons) and anti-NC82 (central complex, (magenta) from flies. The first column is a representative merged image (NC82/GFP); the second column is GFP-stained γ-lobe neurons (green); and the third column is an NC82-stained central complex (magenta). Scale bars, 50µm. **(C)** Quantification of γ lobes via GFP labeling intensity on Day 1 in wildtype controls or *Arc1*^OE^ fly brains. Based on brains displayed in 4D. Asterisks (*) indicate results that are statistically significant at *p-value < 0.05; **p<0.01, ***p<0.001, ****p<0.0001. NS indicates results that show no statistical significance when compared. **(D)** Confocal slices of *Arc1^OE^* and control flies (*yv; attP40*) that express GFP under UAS control in combination with mushroom body driver (*R13F02-Gal4) (R13F02-Gal4>UAS-GFP)* on Day 14. Brains were simultaneously labeled with anti-GFP (mushroom body γ-lobe neurons) and anti-NC82 (central complex, (magenta) from flies. The first column is a representative merged image (NC82/GFP); the second column is GFP-stained γ-lobe neurons (green); and the third column is an NC82-stained central complex (magenta). Scale bars, 50µm. **(E)** Quantification of γ lobes via GFP labeling intensity on Day 14 in wildtype controls or *Arc1*^OE^ fly brains. Based on brains displayed in 4D. Asterisks (*) indicate results that are statistically significant at *p-value < 0.05; **p<0.01, ***p<0.001, ****p<0.0001. NS indicates results that show no statistical significance when compared.

Aged *Rrp40* mutant flies exhibit mushroom body defects (**Figs. 4E-G**) and elevated *Arc1* mRNA levels in each of the three main classes of mushroom body neurons (α/β, α′/β′, and γ) compared to wildtype controls (**Figs. 6B-C**). Thus, we next investigated whether *Arc1* overexpression alone could produce similar mushroom body defects in flies. Accordingly, we assessed the effects of *Arc1* overexpression using the pan-mushroom body driver *R13F02>Gal4* combined with a *UAS-GFP* transgene. On Day 1, overexpression of *Arc1* (*R13F02Gal4/UAS-GFP>UAS-Arc1*) resulted in no obvious mushroom body defects compared to control flies (*R13F02Gal4/UAS-GFP>Ctrl-IR*) via confocal microscopy (**Figs. 8B-C**). Conversely, in aged flies, overexpression of *Arc1* resulted in very thin and faint γ-lobes on Day 14 (**Figs. 8D-E**), reminiscent of mushroom body phenotypes observed in aged *Rrp40* mutant flies. These results suggest that *Arc1* is key in maintaining mushroom body neuronal homeostasis.

## Discussion

This study presents the first cell-type-specific transcriptome analysis of an RNase complex across distinct cell populations within a fully developed animal brain. Specifically, we investigate how the nuclear RNA exosome complex modulates cell-type-specific RNA levels within brain-enriched tissue in *Drosophila* by studying neuropathy-causing RNA exosome variants. By studying pathogenic RNA exosome variants in the fly ortholog of EXOSC3, termed Rrp40, which causes Pontocerebellar Hypoplasia Type 1b (PCH1b) in humans, we aimed to understand how these mutations cause neuronal dysfunction. Utilizing droplet-based snRNA-seq profiling, we show that PCH1b-linked variants (G11A/G11A or G146C/G146C) in RNA exosome structural subunit Rrp40 alter the activity of the complex, causing widespread transcriptomic dysregulation across the 25 distinct cell clusters annotated within fly head tissue, including stabilization of functionally important neuronal transcripts. Notably, we observed aberrant accumulation and distribution of synaptic regulator *Arc1* mRNA across all neuronal and glial subtypes in *Rrp40* mutant brains. Moreover, using long-read Nanopore-based RNA-seq, we show rRNA biogenesis defects in *Rrp40* mutant fly heads. In addition to our molecular analyses, we observed age-dependent morphological defects in the memory-related mushroom body, reduced locomotor ability, and learning and memory impairments in *Rrp40* mutant flies compared to control flies. Importantly, organismal phenotypes in *Rrp40* mutant flies both align with reported neurodegenerative effects in flies (Lu and Vogel, 2009) and PCH1b clinical severity. Crucially, we also show that pan-neuronal overexpression of *Arc1* in aged fly brains phenocopies mushroom body defects observed in *Rrp40* mutant flies. Taken together, our comprehensive analysis of disease-causing PCH1b mutations modeled in *Drosophila* reveals that RNA exosome cap subunit Rrp40 is required to maintain accurate RNA levels and the biogenesis of rRNA across all cell populations within the fly brain and is, consequently, critical for neuronal homeostasis.

Accurately balancing RNA processing and degradation within the cell is achieved by evolutionarily conserved and essential ribonucleases, such as the RNA exosome (Doma and Parker, 2007; West et al., 2004; Zinder and Lima, 2017). Recent reports linking ubiquitously expressed RNA processing factors, including RNases, to tissue-specific neurological disorders (Nussbacher et al., 2019) are confounding, given their essential housekeeping role in intracellular RNA metabolism across all cell types. This tissue-specific pathology raises the question of whether RNases have an enhanced requirement for RNA homeostasis within neuronal cell types. Unbiased transcriptome-wide investigations aimed at deciphering the biological functions of the RNA exosome have been carried out by us and other groups in several biological contexts (Han et al., 2024; Morton et al., 2020; Schneider et al., 2012; Sterrett et al., 2021). Specifically, we performed bulk RNA-seq in fly heads modeling *EXOSC3*/*Rrp40* pathogenic mutations, which revealed aberrant accumulation of functionally important neuronal transcripts, including *Arc1,* other coding RNAs, and non-coding RNAs (Morton et al., 2020). In addition, recent work by another group demonstrates that RNA exosome component 10 (EXOSC10) is critically important for remodeling the transcriptome during key growth-to-maturation events in mouse oocytes (Wu and Dean, 2020). Lastly, an *in vivo* study in yeast confirmed that the RNA exosome interacts with several of its well-known targets, including rRNA, tRNAs, snoRNAs, and snRNAs, as demonstrated through high-throughput PAR-CLiP studies (Sohrabi-Jahromi et al., 2019).

Given that defects in the RNA exosome lead to tissue-specific pathology within the nervous system (Boczonadi et al., 2014; Burns et al., 2018; Di Donato et al., 2016; Morton et al., 2018; Wan et al., 2012), it is critically important to understand how the RNA exosome integrates into cell-type-specific gene expression programs to maintain transcriptome homeostasis. To address this, we employed snRNA-seq to examine nuclear RNA exosome function across all cell populations within head tissue from adult flies modeling PCH1b-linked variants. Intriguingly, our snRNA-seq experiments reveal distinct cell-type-specific molecular signatures within brain-enriched tissue of *Rrp40* mutants (G11A/G11A or G146C/G146C) modeling different PCH1b genotypes (**Fig. 1**). Thus, our transcriptomic analyses support clinical reports of genotype-phenotype correlations in individuals with PCH1b (Rudnik-Schoneborn et al., 2013). Surprisingly, we observed broad transcriptomic dysregulation across all cell populations within *Rrp40* mutant (G11A/G11A or G146C/G146C) head tissue (**Fig. 2**). This result was unexpected, as we initially hypothesized that specific cell types, particularly neurons given the associated clinical phenotypes, would exhibit more significant transcriptome dysregulation compared to non-neuronal cell types. However, we did observe aberrant accumulation and improper distribution of functionally important neuronal transcripts within the fly brain (**Fig. 2B, E**). Intriguingly, the severe PCH1b genotype (G11A/G11A) modeled in flies showed greater transcriptomic dysregulation across brain-enriched cell populations than the mild PCH1b genotype (G146C/G146C). These results are in line with our previous bulk RNA-seq study showing increased transcriptomic dysregulation in G11A/G11A mutants, including *Arc1* mRNA stabilization compared to a mild PCH1b genotype D87A/D87A modeled in *Rrp40* in flies (Morton et al., 2020). These findings provide molecular evidence that distinct pathogenic RNA exosome mutations differentially impact RNA exosome function within brain-enriched tissue. Together, our results demonstrate that pathogenic RNA exosome mutations disrupt the function of the complex, leading to a loss of transcriptome homeostasis across various cell types within the brain.

Consistent with the primary role of the RNA exosome in RNA surveillance (Gudipati et al., 2012; Zinder and Lima, 2017), the stabilization of transcripts in *Rrp40* mutant flies likely results from impaired RNA exosome activity on RNA targets, including specific neuronal transcripts. Whether the transcriptomic dysregulation observed across all cell populations in *Rrp40* mutant fly head tissue is due to defective RNA decay, the accumulation of aberrantly processed transcripts, or both remains to be explored. Further investigation is needed to identify direct targets of the RNA exosome and determine whether the complex preferentially targets specific RNAs, such as *Arc1*, and whether this targeting is cell-type-specific. Developing *in vivo* reporter assays for the RNA exosome would be particularly valuable for understanding how the complex selectively regulates specific RNAs in specific tissues.

In our snRNA-seq experiment on head tissue from *Rrp40* mutant and control flies, we achieved sufficient sequencing depth and power to successfully subcluster and identify neuronal and non-neuronal cell types (**Fig. 1I**), including glial cells, within the fly brain. However, unlike the mammalian brain, where glial cells comprise a substantial portion of the cells within the brain, the *Drosophila* brain predominantly comprises neuronal cells (∼90%) (Raji and Potter, 2021), with a smaller population of glial cells (∼10%) (Freeman, 2015). Notably, there is limited research defining functionally important glial transcripts in *Drosophila,* especially compared to the extensive studies of neuronal transcripts. This fact hindered our efforts to understand how pathogenic RNA exosome mutations affect transcripts important for glial function using our snRNA-seq dataset.

Subcellular compartment-specific cofactors and adaptors guide RNA exosome function and mediate its decision to either process or degrade RNA substrates. How the RNA exosome precisely mediates this dual function remains an area of intense investigation (Zinder and Lima, 2017), and whether specific protein cofactors guide the function of the complex in distinct tissues is unknown. While the functions of the nuclear and cytoplasmic RNA exosome have been elucidated through elegant biochemical studies both *in vitro* and in yeast (Allmang et al., 1999a; Fasken et al., 2017; Gillespie et al., 2017; Mitchell et al., 1997; Schneider et al., 2012) , much remains to be understood about RNA exosome biological functions. The nuclear RNA exosome plays a critical role in rRNA biogenesis by precisely trimming the 3’ ends of specific rRNA species within the nucleolus

Higginson *et al*. 23 (Allmang et al., 1999a; Mitchell et al., 1997). RNA exosome-mediated rRNA biogenesis is conserved across eukaryotes, which is likely why this complex is essential. To investigate rRNA biogenesis in *Rrp40* mutants, we performed northern blotting and long-read Nanopore-based RNA-seq on brain-enriched tissue. We observed 5.8S rRNA biogenesis defects, including accumulation of 5.8S rRNA intermediates and defective degradation of 2S-ITS2 cleavage products (**Fig. 3**). Notably, in a prior study, rRNA processing defects were not detected in fibroblasts from an individual harboring a homozygous D132A amino acid substitution in EXOSC3 which causes a mild form of PCH1b (Wan et al., 2012). These results raise the question of whether distinct RNA exosome mutations differentially alter the function of the complex in rRNA biogenesis or have cell-type-specific consequences depending on the variant.

Maintaining neuronal function relies on the precise regulation of gene expression, including homeostatic mechanisms that balance cellular RNA levels (Bhat et al., 2022; Croset et al., 2018). This balance is particularly critical in neurons, which are highly metabolically active and morphologically complex (Landinez-Macias and Urwyler, 2021). Disruption of RNA-regulatory factors that maintain intracellular RNA homeostasis is increasingly recognized as a key molecular driver of neurological disease (Linder et al., 2015). Here, we show that transcriptome dysregulation caused by disruption of the RNA exosome leads to neurodegenerative effects in *Drosophila*. Specifically, we demonstrate that the brains of *Rrp40* mutants show signs of dysfunctional homeostasis with age, resulting in a significantly increased number of apoptotic neurons within the mushroom body Kenyon cells (**Fig. 5**), which aligns with progressive thinning of mushroom body γ-lobes (**Figs. 4**). These data indicate that Rrp40 is essential for mushroom body neuronal homeostasis. The mushroom body in flies is responsible for olfactory-associated learning and memory (Akalal et al., 2006; de Belle and Heisenberg, 1994; McBride et al., 1999). Given the ubiquitous role of the RNA exosome in maintaining transcriptome homeostasis across all cell populations in the brain (**Fig. 2B**), it is intriguing that specific tissues within the fly brain are more susceptible to transcriptomic dysregulation while others appear more resistant (**Fig. 4**). These findings suggest that the RNA exosome has enhanced or specialized functions in specific cell types. Since we studied *Rrp40* mutant flies during adulthood and aging, we reason that the *Rrp40* mutant phenotypes observed are likely due to cell loss, as neurogenesis is minimal in the adult fly brain (Li and Hidalgo, 2020; Siegrist et al., 2010).

Accurate synaptic transmission is crucial for neuronal communication and function, enabling motor control, learning, and the storage of memories (Hulse et al., 2021; Kennedy, 2013; Rekling et al., 2000). Higginson *et al*. 24 Acetylcholine (ACh), a neurotransmitter produced by cholinergic neurons, plays a key role in regulating these biological processes (Picciotto et al., 2012; Xiao et al., 2016). Intriguingly, analysis of our snRNA-seq dataset reveals that *Rrp40* mutant flies have elevated ChAT levels compared to controls (**Fig. 7D-G**), suggesting that the locomotor impairments observed in G11A/G11A and G146C/G146C mutants (**Fig. 7B**), along with the learning and memory deficits in G11A/G11A mutants (**Fig. 7C**), may be associated not only with mushroom body defects but also with cholinergic dysregulation. Fly models of cholinergic dysfunction show reduced lifespan, impaired locomotor ability, and learning and memory deficits (Showell et al., 2020; Teber et al., 2004; White et al., 2020). Indeed, the phenotypes observed in *Rrp40* mutants in this study mirror reported phenotypes associated with cholinergic dysfunction in flies. In humans, cholinergic synaptic transmission is critical for synaptic plasticity and cell survival in the central nervous system (Resende and Adhikari, 2009; Teber et al., 2004), and disruptions in cholinergic signaling are a hallmark of neurodegenerative diseases such as Alzheimer’s (Chen et al., 2022), Parkinson’s (Muller and Bohnen, 2013), and Huntington’s (Smith et al., 2006).

Protein cofactors and non-coding small RNAs regulate cholinergic signaling at multiple levels (Kratsios et al., 2011; Madrer and Soreq, 2020; Winek et al., 2021). Notably, an essential function of the RNA exosome is the surveillance of non-coding RNAs (ncRNAs) (Allmang et al., 1999a; van Hoof et al., 2000). While our 3’ end poly(A)+ snRNA-seq dataset provides a comprehensive view of coding transcripts, it does not capture the full spectrum of ncRNAs, limiting our ability to draw definitive conclusions about their expression. Therefore, additional high-throughput studies are needed to investigate cell-type-specific regulation of ncRNAs by the RNA exosome. An attractive hypothesis is that the RNA exosome modulates ncRNAs essential for proper cholinergic signaling, thereby supporting neuronal homeostasis and function.

In our snRNA-seq study, we observed significant increases of synaptic regulator *Arc1* transcripts in head tissue of *Rrp40* mutants compared to control flies, with *Arc1* being expressed in 80% of cells in G11A/G11A mutants and 40% in G146C/G146C mutants, compared to about 5% in control flies, where *Arc1* expression is typically restricted to the central brain (**Fig. 2F-G**). These findings are consistent with our previous work reporting *Arc1* accumulation in G11A/G11A mutants (Morton et al., 2020). Intriguingly, recent studies in *Drosophila* models of tauopathy report significant increases in *Arc1* mRNA steady-state levels in the brains of these flies (Zuniga et al., 2023). Furthermore, studies in human neuroblastoma cells show that cholinergic signaling induces *Arc* expression (Soule et al., 2012), thus suggesting that increased *Arc1* mRNA levels observed in *Rrp40* mutant Higginson *et al*. 25 flies may result from a combination of cholinergic signaling dysfunction and impaired RNA exosome function. Together, these data support the hypothesis that increased *Arc1* mRNA steady-state levels could be a molecular driver of neurological disease.

Strikingly, pan-neuronal overexpression of *Arc1* results in increased Arc1 steady-state protein levels in aged flies, causing mushroom body γ-lobe thinning (**Fig. 8D**); thereby, phenocopying observed *Rrp40* mutant mushroom body defects (**Fig. 4E**). This result suggests the RNA exosome is required to tightly regulate *Arc1* mRNA levels to ensure proper brain development and homeostasis. Interestingly, Arc1 steady-state protein levels in *Rrp40* mutants are decreased compared to wildtype control flies (**S5 Fig**) despite *Arc1* mRNA accumulation. This result implies translation efficiency may be impaired in *Rrp40* mutants, which aligns with rRNA biogenesis defects observed in mutant flies (**Fig. 3**) and a recent finding showing that pathogenic RNA exosome mutations modeled in yeast lead to rRNA defects, ultimately reducing the pool of actively translating ribosomes (Fasken et al., 2024). Intriguingly, a recent study reported increased Arc1 protein levels in the adult brains of flies modeling tauopathy (Schulz et al., 2023). This report, along with our findings, suggests that the dysregulation of Arc1 protein levels may serve as a driver of neuronal dysfunction within the fly brain. Reduced Arc1 protein levels observed in *Rrp40* mutant flies suggest that the mushroom body defects observed in these flies likely result from a combination of RNA exosome-mediated transcriptome dysregulation, including increased *Arc1* mRNA steady-state levels and impaired translation efficiency.

In this study, we modeled two patient genotypes associated with PCH1b in *Drosophila* to evaluate the functional consequences of pathogenic RNA exosome structural cap subunit *Rrp40* mutations in the fly brain and investigate the biological function of the RNA exosome in brain-enriched tissue *in vivo*. Our findings reveal that RNA exosome impairment leads to widespread dysregulation of cell-type-specific transcriptomes across the fly brain, with significant dysregulation of functionally important neuronal transcript levels, such as *Arc1.* In addition to transcriptomic analyses, we observed a range of neurodegenerative effects, including mushroom body defects and behavioral phenotypes that support our hypothesis that post-transcriptional gene regulation by the RNA exosome is required for brain homeostasis and function. Notably, although we find that the RNA exosome function is required for controlling RNA abundance across cell populations within the brain, we observe that specific cell types, particularly within the mushroom body, are hypersensitive to RNA exosome dysfunction. These data indicate an enhanced role for the RNA exosome in specific cell types within the fly brain, thus providing a rationale for tissue-specific pathology found in individuals with PCH1b. In summary, our results demonstrate that the RNA exosome cap subunit (EXOSC3/Rrp40) is essential for precisely regulating cellular RNA levels in brain-enriched tissue to maintain neuronal homeostasis and that modeling pathogenic RNA exosome mutations in flies provides valuable insights into PCH1b pathogenesis.

## Materials and Methods

### Drosophila stocks

Crosses were maintained in standard conditions in 25C incubators. Stocks used in this study: *R13*>Gal4, elav- GAL4, UAS-GFP, and the RNA knockdown lines: Control TRiP RNAi (BDSC#36304), *UAS-Rrp40^RNAi^ (TRiP HMJ23923, BDSC #62834)* were obtained from the Bloomington *Drosophila* Stock Center (Indiana, US). *Rrp40^G11A^* and *Rrp40^G146C^* were generated by CRISPR/Cas9 editing at Bestgene, Inc. (CA).

### Generation of CRISPR/Cas9 fly stocks

*Rrp40* mutants and wildtype controls were generated using CRISPR/Cas9 gene editing as previously described (Morton et al., 2020). *pU6-gRNAs*: Target-specific sequences were identified sequences using DRSC fly CRISPR optimal target finder (https://www.flyrnai.org/crispr/) for *Rrp40*. HDR donor vectors (Emory Integrated Genomics Core) were constructed by cloning a 3kb fragment of the *Rrp40* locus, including a loxP-flanked *3xP3-DsRed* cassette inserted downstream of the *Rrp40 3’UTR*, into *KpnI/SmaI* sites of the *pBlueScript-II* vector. Base changes corresponding to G11A and G146C were engineered into this backbone. The *3xP3-DsRed* cassette allows positive selection based on red fluorescence in the adult eye. Injection of DNA mixtures (500 ng/μl HDR and 250 ng/μl U6-gRNA plasmid) into *nos-Cas9* embryos and subsequent screening for dsRed+ transformants was performed by Bestgene, Inc. CRISPR. Newly edited CRIPSR flies were confirmed by Sanger Sequencing at the *Rrp40* locus.

### Single-nuclei RNA sequencing

#### Fly head preparation and nuclei dissociation

Two biological replicates of newly eclosed adult female flies were prepared for each biological replicate ( *Rrp40*^WT^, *Rrp40*^G11A^, and *Rrp40*^G146C^). For each sample, 20 fly heads were pooled together, and nuclei were dissociated using the methods adopted from (McLaughlin et al., 2022; Teefy et al., 2022). For the nuclei dissociation, nuclei were dissociated in 1000µL of homogenization buffer (nuclease free water, sucrose, 10mM Tris (pH 8.0), 25 mM KCl, 5 mM MgCl_2_, 0.1% Triton-x 100, RNasin Plus, 1x protease inhibitor, and 0.1mM DTT) with a 1mL Dounce homogenizer. Homogenization was performed on ice with 20 passes of the loose Dounce and 40 passes of the tight Dounce pestle, and then filtered the solution with a 70µm cell filter into a 15mL conical precoated in 5% Bovine serum albumin. 4mL of nuclei wash buffer (0.5% BSA, PBS, and RNasin Plus) was added to the sample over the filter. Sample was centrifuged, and supernatant was discarded. Another wash step was performed with 5 mL of wash buffer and centrifuged again. Discard the supernatant, resuspend the pellet with 1mL of wash buffer, then transfer to a FACS tube. Add 600µL of debris removal solution (Miltenyi Biotec, Cat# 30-109-398), then 2mL of the resuspension buffer (0.5% BSA, PBS, RNasin) slowly on top. Samples were centrifuged and washed again as described above. Following supernatant removal, the nuclei were resuspended in 150µL of resuspension buffer using a wide-bore tip. The sample was then drawn with a standard pipette and filtered once more using a 40µm Flowmi strainer. Nuclei quantity and quality were measured using a propidium iodine stain visualized by a Flow Cytometer. Only samples with a % debris quantity of 30 or less were utilized for library preparations.

#### 10X genomics single-nuclei library preparation

Single-cell libraries were prepared using Chromium Next GEM Single Cell 3ʹ GEM, Library & Gel Bead Kit v3.1 (10X Genomics, PN- 1000213) according to manufacturer’s instructions (10x Genomics User Guide Chromium Next GEM Single-cell 30 Reagent Kits v3.1 (CG000204, Rev D)). Based on cell estimates from flow cytometry, cell suspensions were loaded for a targeted cell recovery of 10,000 cells per sample. The prepped libraries were quantified on a Qubit 3.0 Fluorometer (Thermo Fisher Scientific, Q33216). Completed single-cell libraries were assessed for quality on the 4200 TapeStation system (Agilent Technologies; G2991A) with a High Sensitivity D1000 DNA ScreenTape (Agilent Technologies 50675584). Libraries were sequenced on an Illumina Novaseq 6000 generating 150 bp paired-end reads at Novogene USA.

#### Data processing and computational analysis

The reads were mapped to BDGP *D*. *melanogaster* (r6.25) genome. Cell ranger version 7.1.0 (10X Genomics) (Zheng et al., 2017) was used to create the reference. Quality control of the data was performed in R version 4.1.1 using Seurat version 4.2.1 (Butler et al., 2018; Hao et al., 2024; Stuart et al., 2019) by removing dead and low-quality cells and selecting cells containing between 300 and 4000 UMIs and <10% mitochondrial reads. Following quality control, all cells from each genotype were pulled together and normalized using SCTransform version 0.3.5. Following normalization, PCA and 2D UMAP dimension reduction analyses were performed. UMAP analysis was conducted with 30 dimensions and a resolution of 5. Manual annotation of the clusters was performed using published cellular markers from (Davie et al., 2018; Li et al., 2022). Following annotation of distinct cell types, cells were re-classified based on neuronal, glial, head-specific, and unannotated. Pseudobulk analysis was performed with the Muscat package version 1.8.2 and DESeq2 version 1.34.0. In addition, subclustering analysis was performed on all cell types, including Kenyon cells, using the subset function based on cell identities. Further, differential expression analysis was also confirmed using the 10X genomics platform Loupe Browser 8. An UpSet plot was created in R version 4.1.1 to compare the overlap number of transcripts elevated in *Rrp40* mutants (G11A/G11A and G146C/G146C) neuronal and glial cells (log2FC>2). Biological processes linked to the overlapping genes were identified using the FlyEnrichr database (Chen et al., 2013; Kuleshov et al., 2019; Kuleshov et al., 2016).

#### Hybridization Chain Reaction RNA-Fluorescent *in situ* Hybridization + IHC

This protocol was modified from the Molecular Instruments HCR™ IF + HCR™ RNA-FISH protocol for sample in solutions. Brain dissections and staining were performed as described previously(Morton et al., 2020; Wu and Luo, 2006). Brains of anesthetized flies were dissected in 1x PBS, then transferred into PBS-tween (0.2%) and fixed in 4% paraformaldehyde (32% PFA, Electron Microscopy Sciences, Cat #15714-S) overnight at 4C. Following fixation, the brains were washed in PBS-tween and preserved in 100% methanol overnight at 4C. The tissue was then rehydrated and permeabilized with 0.3% PBS-Triton for 20 minutes at room temperature. Following permeabilization, samples were blocked in antibody diluent (Molecular Instruments) for 30 minutes at room temperature and incubated with primary antibody (NC82 anti-bruchpilot obtained from Developmental Studies Hybridoma Bank (DSHB) 1:50) overnight at 4C. After wash steps, samples were incubated with the initiator-labeled secondary antibody (1:500) overnight at 4C. Following wash steps, samples were post-fixed with 4% PFA for 10 minutes at room temperature. After fixation, samples were washed with 5x SSCT and pre-hybridized in the probe hybridization buffer for 15 minutes at 45C. 0.2 pmol of the 1µM DNA probes complementary to *Arc1* mRNA (designed by Molecular instruments) was then added to the solution, and samples were incubated overnight at 45C. Next, samples were washed using Probe Wash Buffer (Molecular Instruments) and 5x SSCT. Following wash steps, samples were blocked with amplification buffer at room temperature for 15 minutes. 2µL of each activated (snap-cooled) fluorescently labeled hairpins were added to the solution and incubated overnight in the dark. Samples were then washed with 5x SSCT and mounted using Gold antifade mountant and imaged using the Zeiss 880 LSM confocal microscope at 20X magnification. Each image was processed and quantified in Imaris microscopy image analysis software. Quantification of each sample was performed on the mean max intensity of each sample. Statistical analysis was performed in PRISM (GraphPad, San Diego).

### Nanopore Sequencing

#### Library Preparation

1 ug of total RNAs were isolated from approximately 20 fly heads using TRIzol and *in vitro* polyadenylated using E. coli Poly(A) Polymerase (NEB, catalog #M0276S). RNA libraries were prepared from 500 ng of in vitro polyadenylated RNAs using the PCR-cDNA barcoding kit from Oxford Nanopore (ONT, catalog #: SQK- PCB114.24) as per the manufacturer’s instructions. Sequencing was performed using R10.4.1 flongles on a MinION Mk1B device and sequenced for 48 hours. Basecalling was performed using the built in Dorado basecaller in Minknow (Dorado version: 7.4.12). Reads were then mapped to a single copy of the *Drosophila melanogaster* ribosomal RNA primary transcript using Minimap 2 (Version 2.17-r941). Reads were visualized using IGV (Version 2.12.3).

#### Nanoblot Sequencing Quantification

For read counts, we calculated the number of reads mapping to each gene using featureCounts (Liao et al., 2014). DESeq2 was used to quantify changes in gene expression of the different strains and for normalizing counts to library size(Love et al., 2014). Nanoblot was used to visualize amounts of 2S rRNA extensions for 3 samples (DeMario et al., 2023).

### Brain dissections and immunochemistry

#### Mushroom body staining via anti-Fasciclin II antibody

Brain dissections and staining were performed as described previously (Wu and Luo, 2006). Brains of anesthetized flies were dissected in 1x PBS, fixed in 4% paraformaldehyde ( 32% PFA, Electron Microscopy Sciences, Cat #15714-S ) overnight at 4C, washed 3 times in 1 x PBS, permeabilized with 0.3% PBS-Triton, and then stained for 2 days with primary antibody (1D4) diluted in antibody solution (0.1% PBS-Triton and Normal Goat Serum (NGS)). Following several washes with 0.1% PBS-Triton, brains were incubated with the appropriate fluorescently conjugated secondary antibody (1:500) in antibody solution overnight at 4C, washed in 0.1% PBS-Triton, and then mounted in ProLong Gold antifade reagent (Invitrogen). The 1D4 anti-FasII hybridoma (1:20) developed by C. Goodman(Grenningloh et al., 1991) was obtained from the DHSB. Brains were imaged using the Zeiss LSM 880 confocal microscope.

#### Fly brain staining via anti-DLG and anti-GFP antibodies

Brains of anesthetized flies were dissected in ice-cold 0.05% PBS-Triton, fixed in 4% paraformaldehyde overnight at 4C. Following fixation, brains were washed three times in PBST for 1 hour. Following wash steps, brains were stained with primary antibody (NC82) diluted in antibody solution (970µL of PBST, 30µL NGS, 1µL rabbit anti-GFP (Cell Signaling Technology, Cat # 2956S, 1:1000), 50 µL mouse anti-DLG (1:20)) for 3 hours at RT then overnight at 4C. Samples were washed three times for 1 hour in PBST, then incubated with the appropriate fluorescently conjugated secondary antibody (1:500) in antibody solution for 3 hours at RT, then 3-4 days at 4C. Samples were re-washed three times for 1 hour, then mounted in ProLong Gold antifade reagent on a microscope slide with spacers separating the coverslips to prevent brain tissue damage. Brains were imaged at 40X using the Leica SP8 confocal microscope.

### Behavioral Assays

#### Lifespan analysis

Lifespan was assessed in standard conditions at 25C. Newly eclosed animals were collected, separated by sex, placed into new vials (up to 15 per vial), and transferred to fresh vials weekly. Survivorship was scored daily for all genotypes. At least 100 flies of each genotype were tested. Log-rank (Mantel-Cox) test of survivorship curves was performed using PRISM (GraphPad, San Diego).

#### Locomotor Function Assay

Negative geotaxis assays were performed as previously described(Pak et al., 2011) with some modifications. Newly eclosed *Rrp40* flies (WT/WT, G11A/G11A, G146C/G146C) (day 0) were collected, divided into groups of 12, and kept in separate vials. Cohorts of age-matched flies were then transferred to a 25-ml graduated cylinder to analyze climbing response.

#### Learning and Memory

Tastant-induced proboscis extension learning and memory assays were performed as described(Li et al., 2020; Wang et al., 2004). The proboscis extension response assay measured taste and short-term memory in mutant and wildtype flies. First, flies were starved for 18 hours prior to testing by placing ten flies in a vial with only a Kim-wipe damp with water. Each genotype had six groups of 10 flies. Following the starvation period, 10 flies were placed onto a microscope slide by their wings, with a separate slide for each genotype. To prevent any alterations in behavior, flies were immobilized using ice instead of CO_2_. Once the flies were fixed onto the slide, with only head mobility, we prepared the sucrose Kimwipes. To ensure minimal sucrose contact, we twisted the tip of a Kimwipe into a thin thread before dipping it into 500µM sucrose solution (Sucrose, dH2O) (EMD, Cat #SX1075-1). During the pre-testing period, before providing tastants, we gave the sucrose solution to all the flies to act as a positive control. Any flies that did not extend their probiscis were not used in the study. Following the pretest, we trained the flies by presenting them with a caffeine solution, the deterrent (Caffeine 20µM (Sigma Aldrich Cat #CO750-500G), followed by the sucrose solution, the attractant. The training period was performed at 5-minute intervals for 15 minutes. Following the training period, we tested the flies’ short-term learning and memory capability, by presenting them with the sucrose solution after the training period (0 minutes) and after 30 minutes. Flies with defects in olfactory learning and memory will have a higher proboscis extension response to the sucrose solution following the training period.

#### Immunoblotting

20µg protein lysates of fly heads were resolved on 4–20% Criterion TGX polyacrylamide gels (BioRad), transferred to nitrocellulose membranes, incubated for 5-10 minutes in blocking buffer (Everyblot Blocking Buffer, BioRad #12010020), followed by an overnight incubation at 4°C in primary antibody diluted in blocking buffer. Primary antibodies were detected using species-specific horse radish peroxidase (HRP) conjugated secondary antibodies (Invitrogen) and enhanced chemiluminescence (ECL, Sigma). Primary antibodies include rabbit anti- *Arc1*, which was provided by Dr. Travis Thomson, and mouse anti-GAPDH (1:1000, Proteintech).

## Data Availability Statement

All relevant data are within the manuscript and its Supporting Information files. The data from the snRNA-Seq have been uploaded at NCBI GEO (accession no. GSE280166). All scripts used to analyze this dataset are available on the Morton lab GitHub at https://github.com/themortonlab/flyprojects. Nanopore-based RNA-seq analysis has been uploaded to NCBI BioProject Database (BioProjectID: PRJNA1179429).

## Acknowledgments

This work was supported by a National Institutes of Health R01 grant (NS131620), Alfred P. Sloan Research Fellowship in Neuroscience award (FG-2023-20698), and a University of Southern California-Buck Institute Nathan Shock Center Award (P30AG068345) to D.J.M. L.A.H was supported by the National Science Foundation Graduate Research Fellowship Program and a University of Southern California Provost Fellowship. B.A.B. was supported by a Simons Foundation award (SF811217).

## Supplemental Figure Legends

**SFig. 1:**
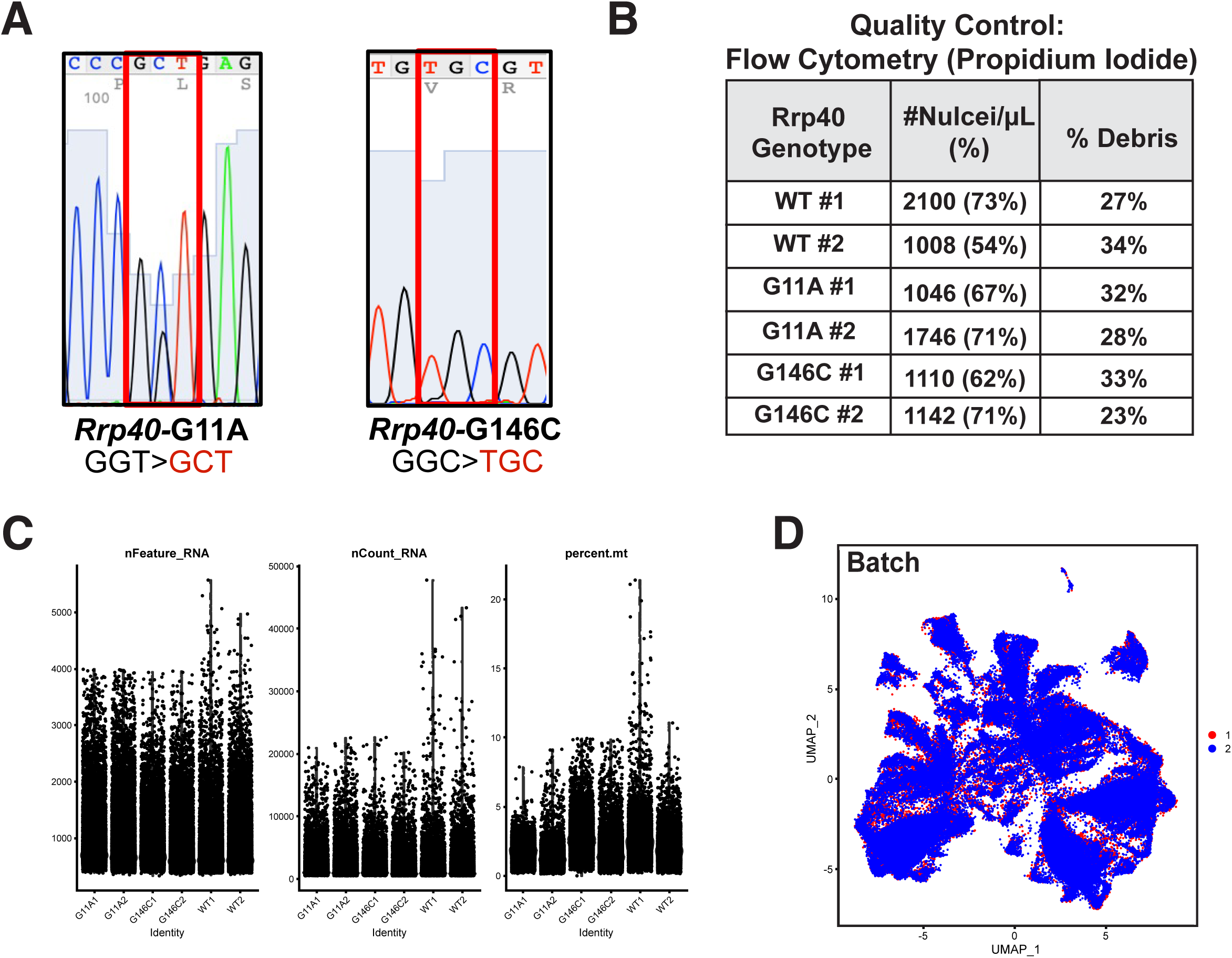
Quality control analysis on single nucleus RNA-sequencing of confirmed CRISPR/Cas9 genetically engineered *Rrp40* mutant and wildtype control flies. **(A)** Electropherogram validating the presence of desired missense mutations in the gene encoding RNA exosome subunit, Rrp40 using Sanger Sequencing. **(B)** Chart depicting the quality of single nuclei isolation by measuring % single nuclei and % cellular debris using Flow Cytometry and propidium iodide staining. **(C)** Violin plot representing the number of nFeatures (genes detected per cell), nCounts (number of unique molecular identifiers per cell), and percent mitochondria in each replicate of the *Rrp40*-WT, G11A/G11A, and G146C/G146C sequencing results. This readout was used to determine the quality of the library preparation and establish cutoffs on the dataset for downstream analysis. Data points with a nFeature_RNA > 300 and nFeature_RNA <4000, percent.mt <10, and log10GenesPerUMI > .80 were kept for single nucleus RNA sequencing analysis. **(D)** 2D UMAP plots of snRNA-seq data from *Rrp40* mutant (G11A/G11A G146C/G146C) and *Rrp40* wildtype control samples by Batch (Batch 1 (red), Batch 2 (blue)).

**SFig. 2:**
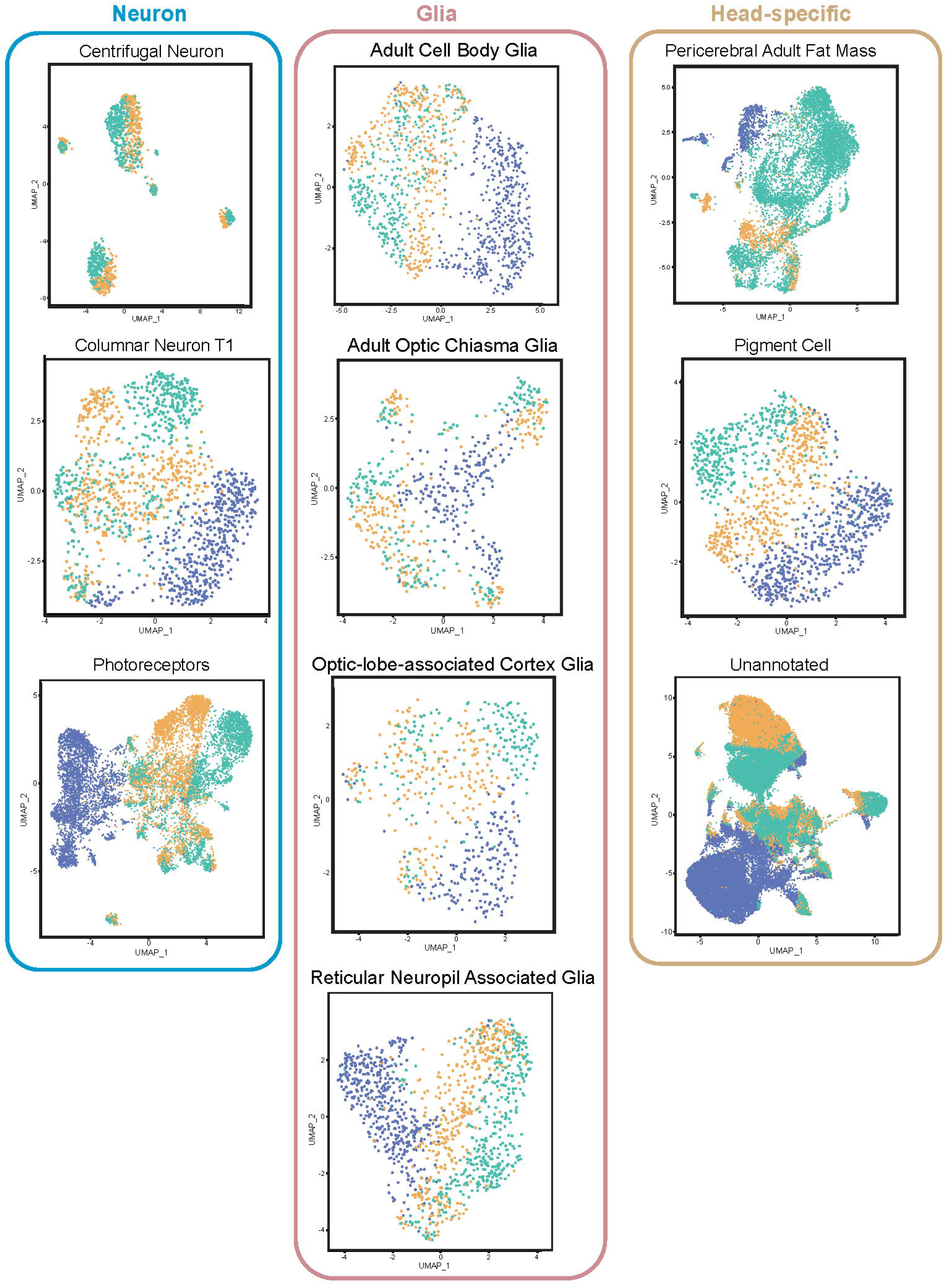
2-Dimensional Reduction Analysis on sub-clustered cell types from neuronal, glial, and head-specific cell classes. Sub-clustered neural, glial, and head-specific cell types visualized by 2D UMAP plots colored by *Rrp40* genotypes (WT/WT (blue), G11A/G11A (red), and G146C/G146C (green).

**SFig. 3:**
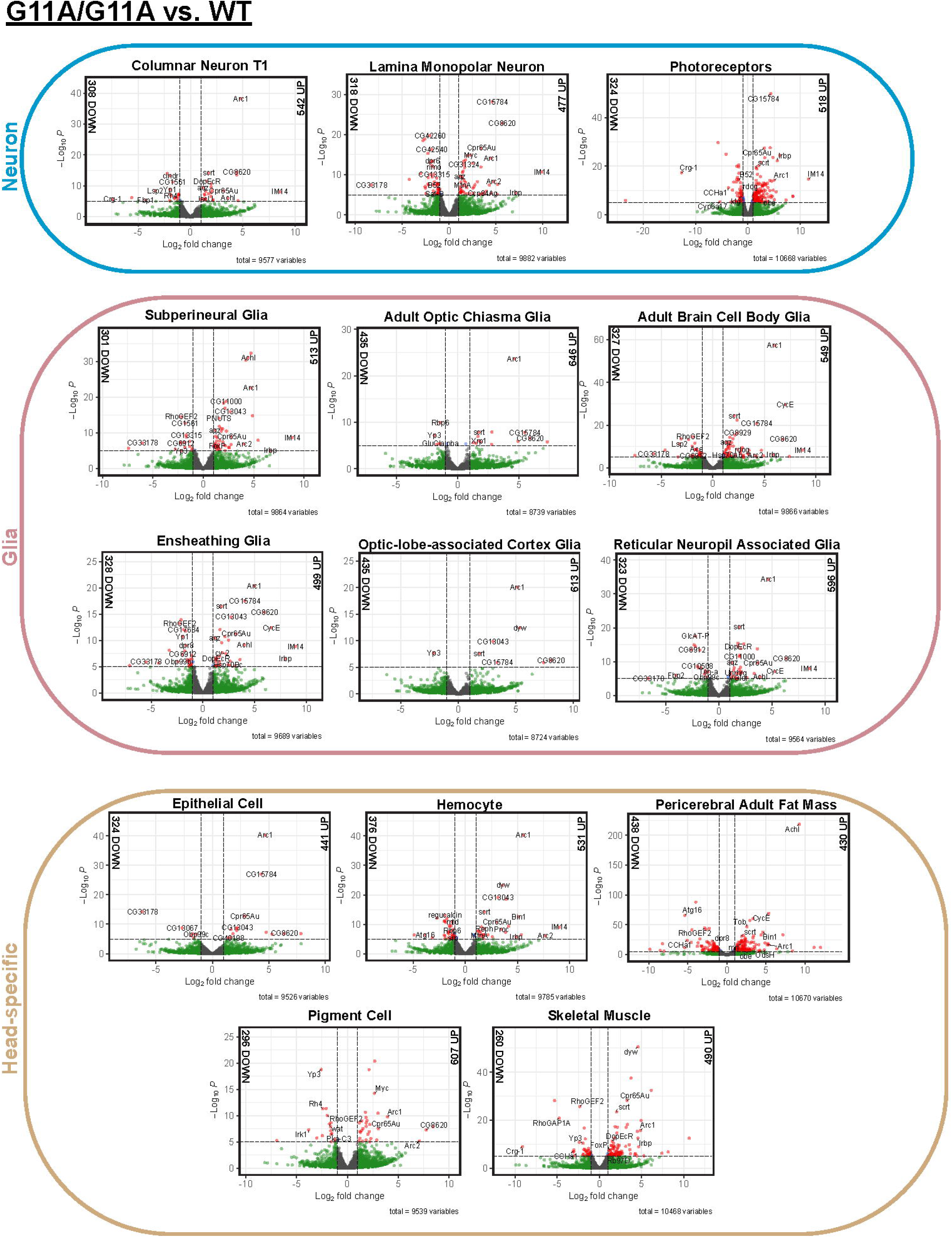
Differential transcript abundance analysis on severe G11A/G11A mutant versus wildtype control. Volcano plot representing differentially expressed transcripts in G11A/G11A vs. WT/WT neuronal, glial, and head-specific cell types. Transcripts (absolute Log2FC > 1) (green), transcripts (p-value < 1e-05) (blue), transcripts (absolute Log2FC > 1 and p-value < 1e-05) (red), and transcripts that meet neither criterion are in gray. The number of transcripts with absolute Log2FC >2 is indicated in black.

**SFig. 4:**
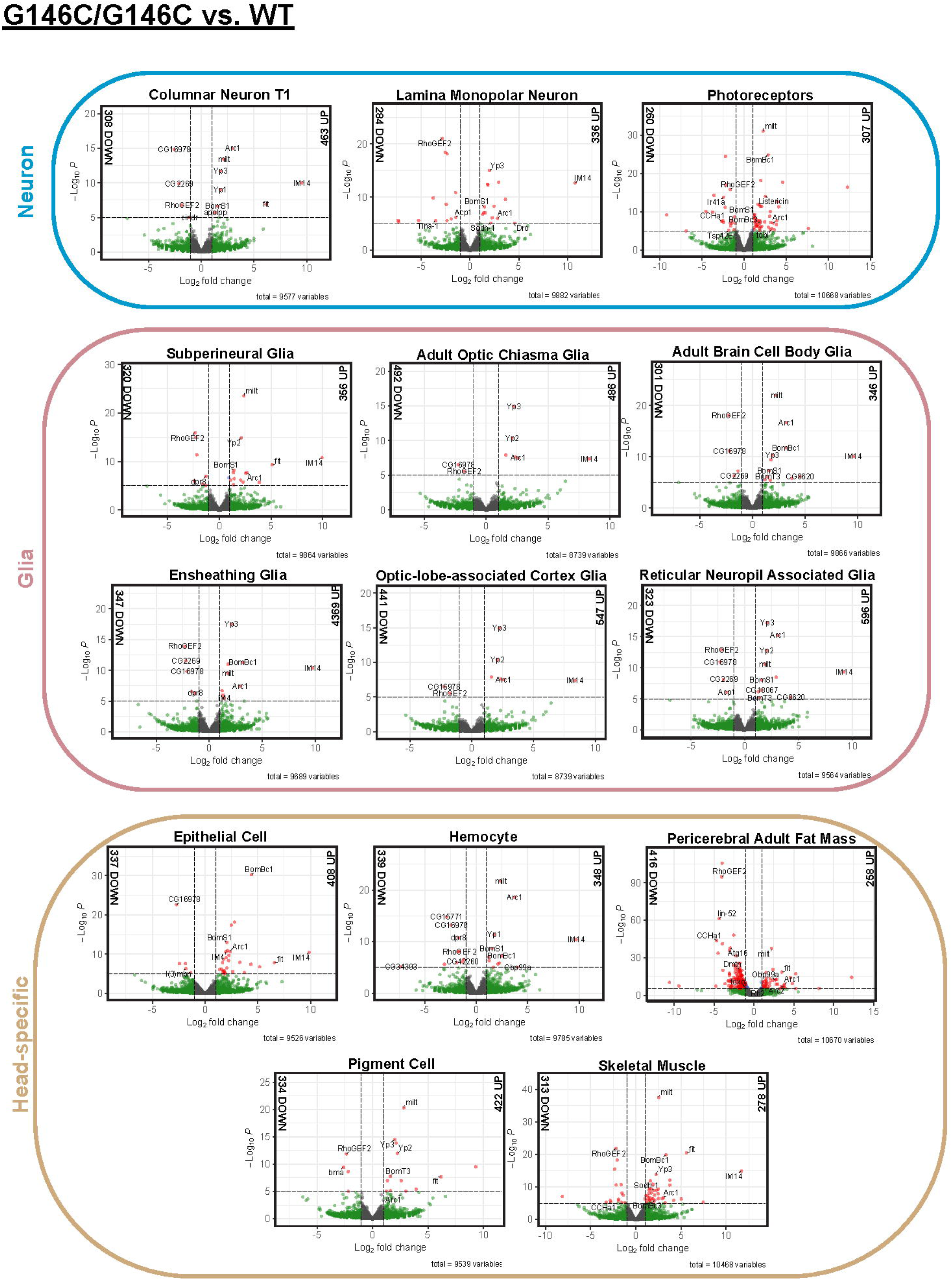
Differential transcript abundance analysis on mild G146C/G146C mutant versus wildtype control. Volcano plot representing differentially expressed transcripts in G146C/G146C vs. WT/WT neuronal, glial, and head-specific cell types. Transcripts (absolute Log2FC > 1) (green), transcripts (p-value < 1e-05) (blue), transcripts (absolute Log2FC > 1 and p-value < 1e-05) (red), and transcripts that meet neither criterion are in gray. The number of transcripts with absolute Log2FC > 2 is indicated in black.

**SFig. 5:**
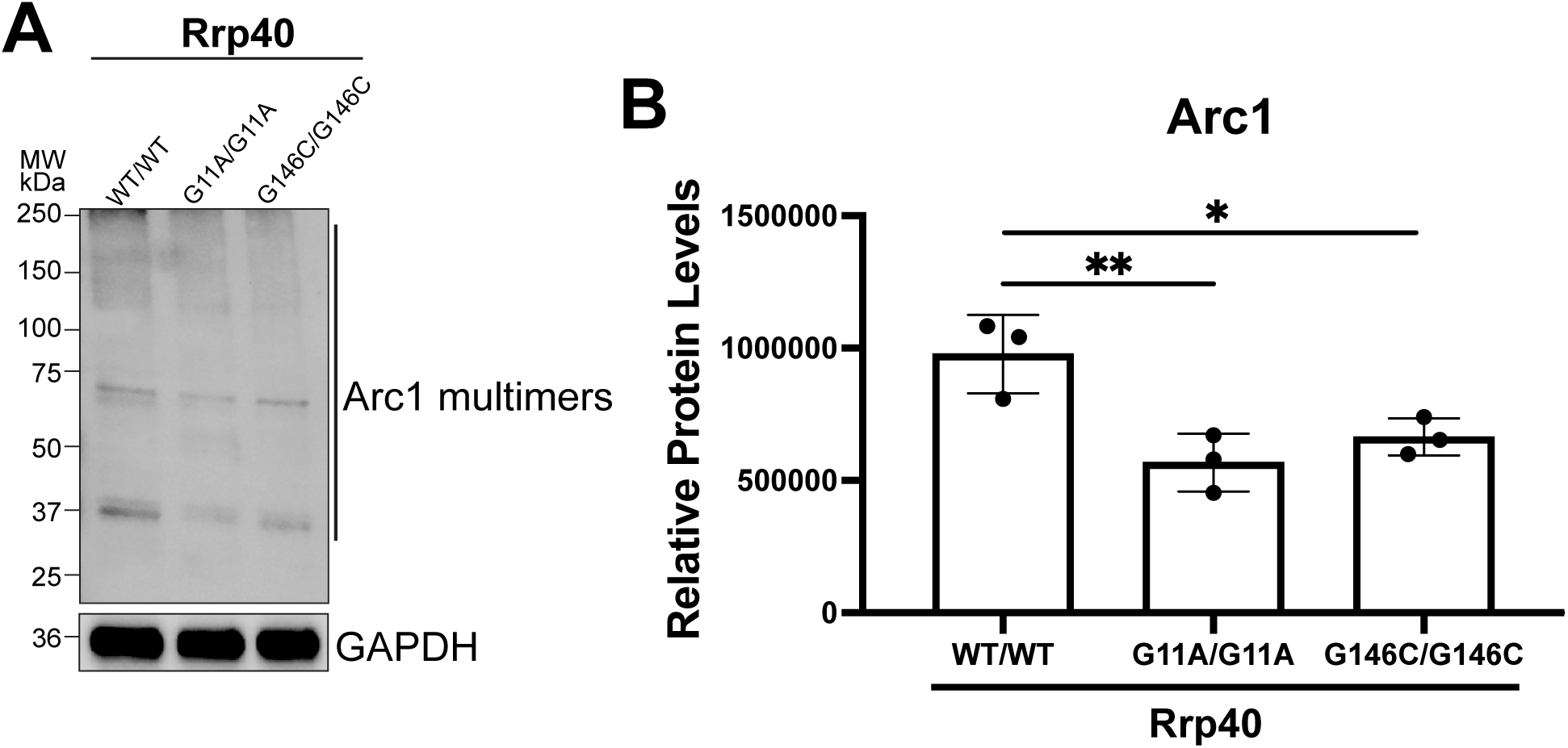
PCH1b causal variants modeled in *Drosophila* alter Arc1 steady-state protein levels. Lysates prepared from heads of control WT/WT or *Rrp40* mutant flies (G11A/G11A or G146C/G146C) were resolved by SDS-PAGE and analyzed by immunoblotting with antibodies to detect Arc1. Both GAPDH and Stain Free, as a measure of total protein, serve as loading controls. A representative immunoblot is shown (n=3). **(B)** Results from (A) were quantified for Arc1 and are presented as Relative Protein Levels with Rrp40 mutants (G11A/G11A or G146C/G146C) compared to control (WT/WT) flies. Asterisks (*) indicate results that show statistical significance at *p-value <0.05 and **p-value<0.01.

**SFig. 6:**
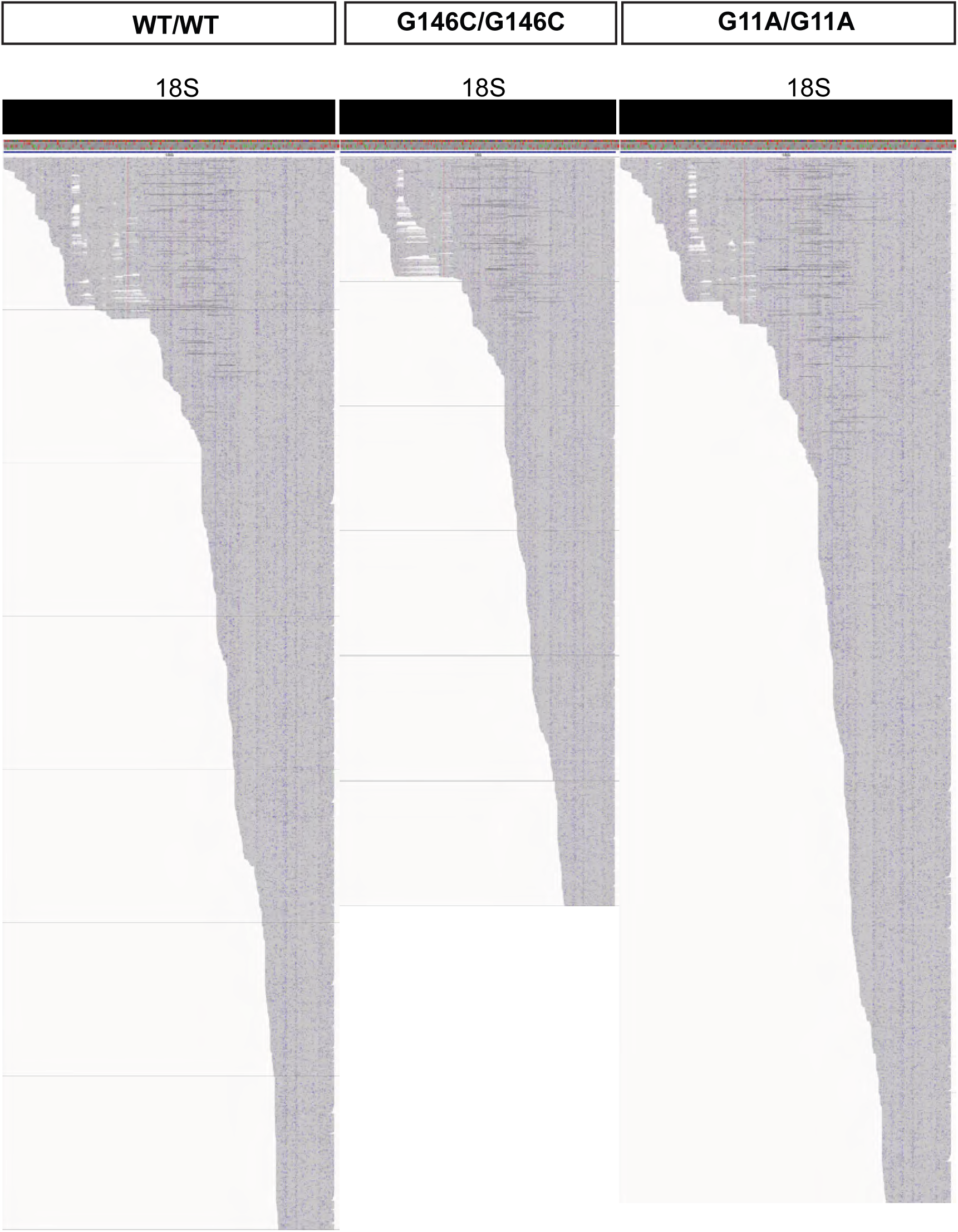
Nanopore long read sequencing in *Rrp40* mutant head tissue reveals no 18S rRNA biogenesis defects. Integrated genome viewer (IGV) screenshot of single-track analysis of reads mapping to 18S rRNA in *Rrp40* mutants (G11A/G11A, G146C/G146C) and wildtype control.

**SFig. 7:**
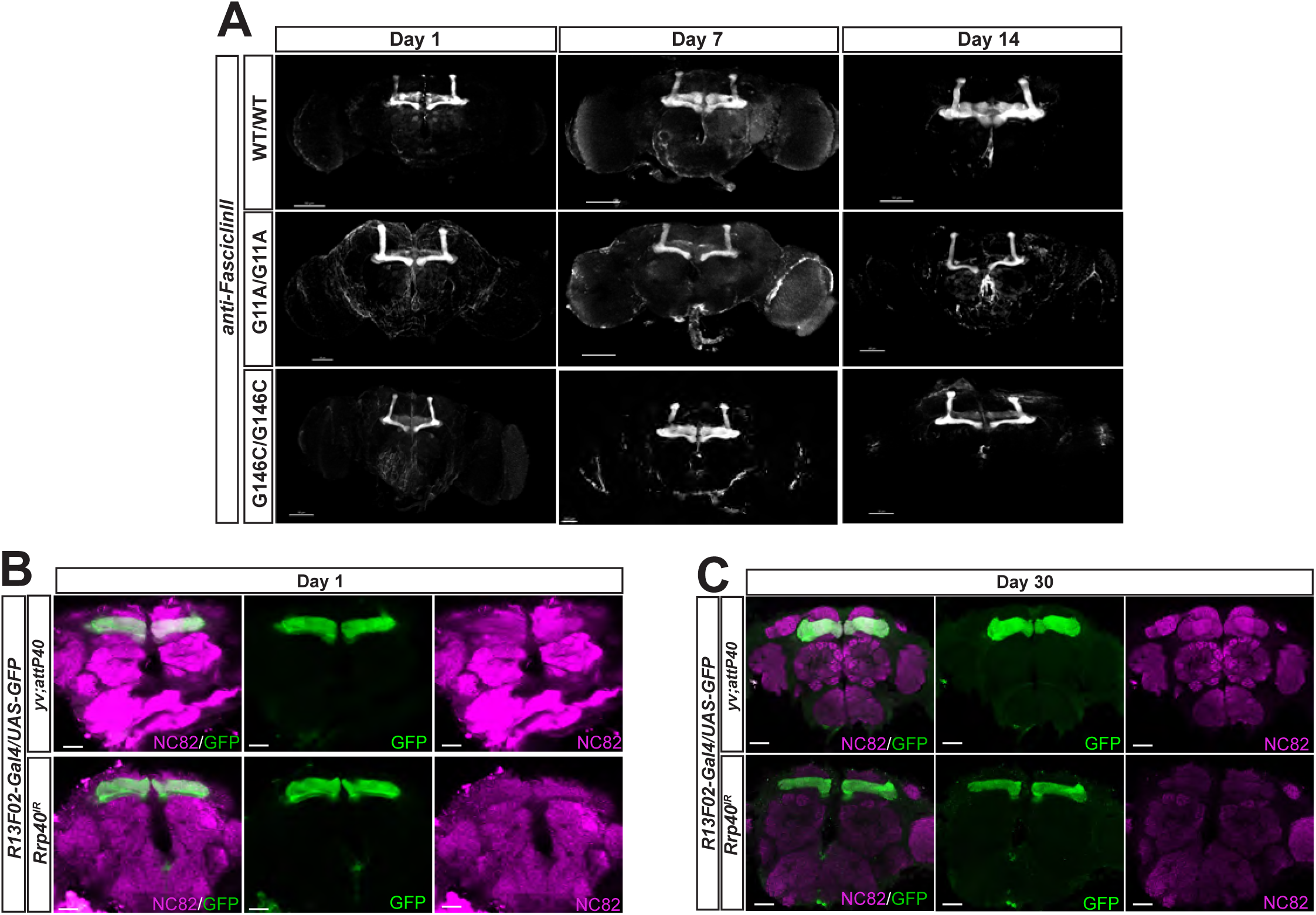
Diverse labeling of the mushroom body brain structure confirms *Rrp40* mutants show age-dependent mushroom body defects. **(A)** Mushroom Body structure stained by anti-Fascillin II (FasII) in *Rrp40* mutant (G11A/G11A or G146C/G146C) and wildtype fly brains (n = 10) at Day 1, 7, and 14 aged adult female flies (scale = 30-50µm). **(B)** Max intensity projection of anti-DLG staining the mushroom body lobes within *Rrp40* mutant and wildtype brains at Day 1 and Day 14 aged adult flies. (n = 8, scale = 50µm) anti-DLG staining was performed to confirm the absence of FasII staining in the mushroom body γ lobes. **(C)** Confocal slices of *Rrp40^IR^* and control flies (*yv; attP40*) that express RNAi and/or GFP under UAS control in combination with mushroom body driver (*R13F02-Gal4) (R13F02-Gal4>UAS-GFP)* on Day 30. Brains were simultaneously labeled with anti-GFP (mushroom body γ-lobe neurons) and anti-NC82 (central complex, (magenta) from flies. The first column is a representative merged image (NC82/GFP); the second column is GFP-stained γ-lobe neurons (green); and the third column is an NC82-stained central complex (magenta). Scale bars, 50µm.

